# A Single Cell Cloning Platform for Gene Edited Functional Murine Hematopoietic Stem Cells

**DOI:** 10.1101/2022.03.23.485423

**Authors:** Hans Jiro Becker, Reiko Ishida, Adam C. Wilkinson, Takaharu Kimura, Michelle Sue Jann Lee, Cevayir Coban, Yasunori Ota, Arinobu Tojo, David Kent, Satoshi Yamazaki

## Abstract

Gene editing using engineered nucleases frequently produces on- and off-target indels in hematopoietic stem cells (HSCs). Gene-edited HSC cultures thus contain genetically heterogenous populations, the majority of which either do not carry the desired edit or harbor unwanted mutations. In consequence, transplanting edited HSCs carries the risks of suboptimal efficiency and of unwanted mutations in the graft. Here, we present an approach for expanding gene-edited HSCs at clonal density, allowing for genetic profiling of individual clones before transplantation. We achieved this by developing a defined, polymer-based expansion system and identifying long-term expanding clones within the CD201^+^CD150^+^CD48^-^c-Kit^+^Sca-1^+^Lin^-^ population of pre-cultured HSCs. Using the *Prkdc*^scid^ immunodeficiency model, we demonstrate that we can expand and profile edited HSC clones to check for desired and unintended modifications. Transplantation of *Prkdc-*corrected HSCs rescued the immunodeficient phenotype. Our *ex vivo*-manipulation platform establishes a novel paradigm to control genetic heterogeneity in HSC gene editing and therapy.

## Introduction

The rapid adoption of engineered nucleases has put hematopoietic stem cells (HSCs) at the center of gene editing applications. The ability to functionally interrogate genes by introducing or correcting mutations at precise loci has greatly advanced our understanding of HSC biology and has enabled curative approaches for genetic diseases. CRISPR/Cas9 currently represents the most widespread system for gene editing of the hematopoietic system (Naldini, 2019). A target-specific guide RNA (gRNA) directs the Cas9 endonuclease to a genomic site of interest, where it induces a DNA double strand break (DSB). The subsequent engagement of the cell-intrinsic DNA damage repair (DDR) machinery can be exploited to create targeted modifications in HSCs (Dever et al., 2016; De Ravin et al., 2017; Schiroli et al., 2017). Since mutagenic repair (e.g. non-homologous end joining (NHEJ)) takes precedence in primitive HSCs (Mohrin et al., 2010), a phenomenon closely tied to their dormant phenotype, random small insertions and deletions (indels) represent the most common on-target editing outcome (Dever et al., 2016; Genovese et al., 2014). In contrast, correction via templated repair (i.e. homology-directed repair (HDR)) among long-term (LT)-HSCs remains inefficient (Lattanzi et al., 2021; Mohrin et al., 2010). Off-target mutations may also raise concerns about genotoxicity in Cas9-edited cells (Tsai and Joung, 2016). Together, these unwanted mutations may confound the effects of the edited gene of interest and represent incalculable risks in basic and translational research settings. Apart from gene editing, maintaining HSC self-renewal in long-term cultures required by gene editing protocols remains challenging (Wilkinson et al., 2020). Consequently, edited and bulk-expanded HSC cultures contain genetically and functionally heterogenous populations and only include a low fraction of functional HSCs with the desired genetic modifications.

Expansion of single, *bona fide* HSCs would overcome this limitation by enabling direct profiling of on- and off-target editing outcomes, allowing for selective transplantation only of clones with a defined mutational pattern. However, current protocols do not allow for expansion of HSCs at clonal density to the extent necessary for transplantation. Clonal expansion technologies for embryonic stem cells (ESCs) and induced pluripotent stem cells (iPSCs) have been a major driver for advances in the biology and translational research of PSCs, yet the generation of functional HSCs from PSCs remains a major hurdle.

We recently reported on a serum-free, polyvinyl alcohol (PVA)-based HSC expansion protocol that permits up to 899-fold expansion of HSCs over a period of 4 weeks (Wilkinson et al., 2019). Here, we use this protocol to show that bulk expansion produces a genetically heterogenous graft with on- and off-target indels. Addressing this issue, we present a novel system that supports single cell expansion of edited HSCs and define a phenotype that assists in selecting precultured clones with long-term expansion potential. We apply this system to a gene correction model of severe combined immunodeficiency (SCID), demonstrating the feasibility of single cell expansion for sequence-based selection of edited HSC clones.

## Results

### Gene-edited HSCs correct Prkdc^scid^ immunodeficiency but bear on- and off-target indels

The immunodeficient phenotype in CB17/SCID mice is caused by a T to A mutation in the *Prkdc* gene (*Prkdc*^scid^), creating a premature termination codon (PTC; p.Y4046X) and leading to functional loss of its product, DNA-dependent protein kinase catalytic subunit (DNA-PKcs, Fig. 1A) (Araki et al., 1997). DNA-PKcs is indispensable for the resolution of DNA double strand breaks (DSBs) during V(D)J recombination, which is reflected in the absence of functional B and T cells in CB17/SCID mice.

**Figure 1.**
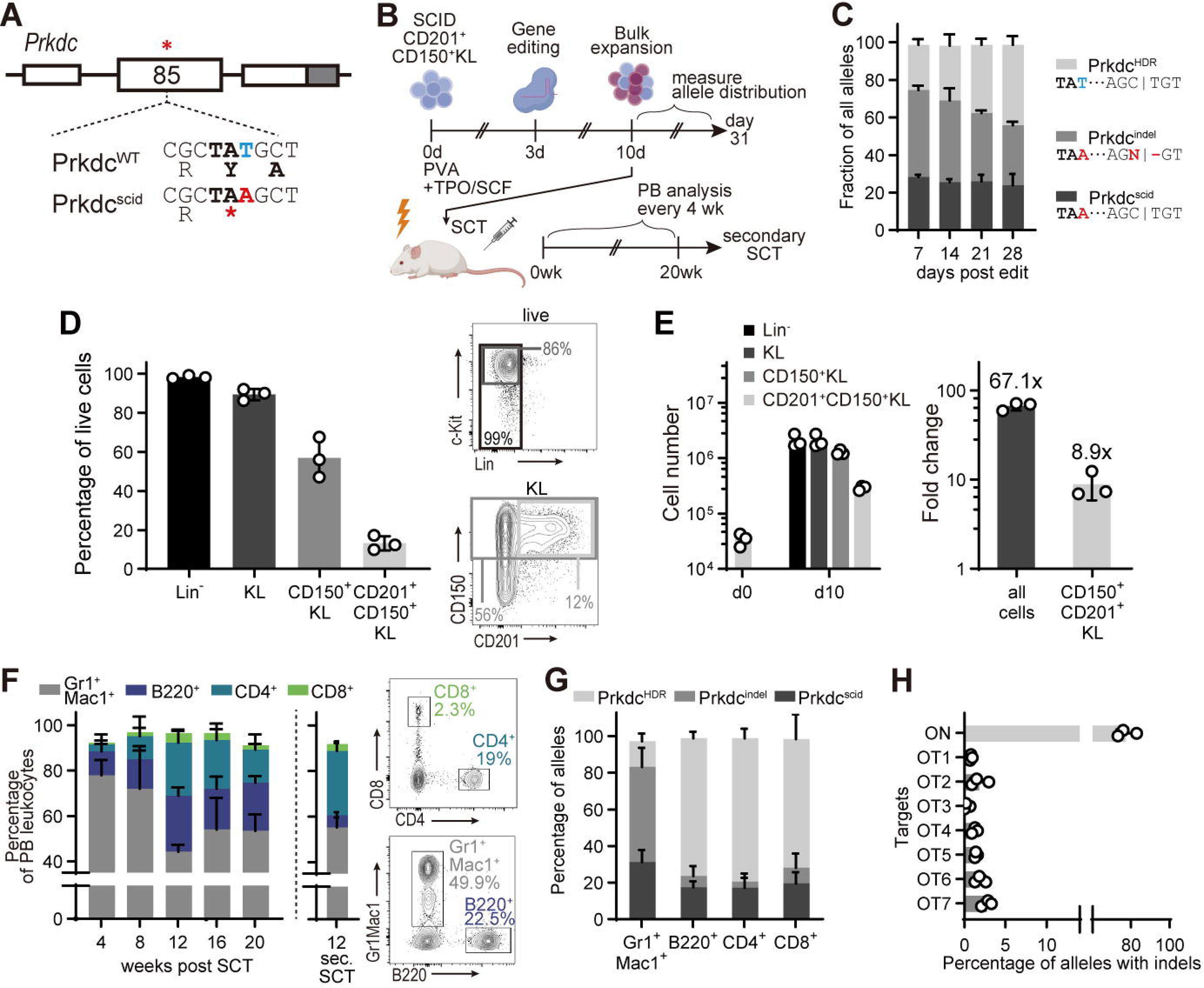
Autologous HSCT gene correction rescues the *Prkdc*^scid^ phenotype but introduces on- and off-target indels. **(A)** Genomic context of the *Prkdc*^scid^ mutation in exon 85. White boxes: exons, grey box, 3’UTR.* denotes location of *Prkdc*^scid^ mutation. **(B)** Experimental scheme of the gene editing and HSC expansion model. **(C)** Post-editing allele distribution at the *Prkdc* locus, assessed by ICE (n=3 cultures). **(D)** Fractions of immunophenotypically defined HSPC populations within cultures on day 10 of culture, 7 days post-editing. Percentage of all live cells (n=3 cultures). **(E)** Absolute cell numbers (left panel) and fold-change expansion (right panel) of cultured HSPCs, day 10 of culture. **(F)** Left: Frequencies of peripheral blood (PB) leukocytes as percentage of all live leukocytes (n=3 groups, 3-4 mice per group). Plot next to dashed line shows frequencies 12 weeks post-secondary SCT (n=5 mice). Right: representative FACS plots 20 weeks post-transplant. **(G)** Frequencies of *Prkdc* alleles in sorted PB cells 20 weeks post-SCT (n=3 experiments, 3-4 mice per group). **(H)** On- and off-target (OT) activity of the *Prkdc*-specific gRNA, assessed with TIDE. The seven highest scoring off-target sites, as predicted by COSMID, were interrogated. See Table S1 for detailed information about the off-target sites interrogated.

To determine whether the *Prkdc*^scid^ phenotype can be corrected with gene edited and bulk-expanded HSCs and to assess the levels of indels generated in the process, we designed a gene editing protocol based on our previously established *ex vivo* HSC expansion platform (Fig. 1B) (Wilkinson et al., 2019). CD201^+^CD150^+^c-Kit^+^Lin^-^ cells from CB17/SCID mice were cultured in PVA-based medium (PVA-HSC) for three days (Fig. S1A). We included CD201 (EPCR) in our isolation panel since the commonly employed marker Stem Cell Antigen 1 (Sca-1) is known to be poorly expressed on hematopoietic cells of non-C57BL/6 mouse strains and because CD201 has shown to be a reliable marker in BALB/c mice, from which the CB17/SCID strain is derived (Vazquez et al., 2015). Cas9 ribonucleoprotein (RNP) complexes and a corrective HDR template were delivered into HSCs three days after isolation. One week after editing, the majority of alleles contained indels (*Prkdc*^indel^, 48%), while 26% had incorporated the HDR donor sequence (*Prkdc*^HDR^, Fig. 1C). HDR frequencies were lowest (11±2%) in the most stringently defined HSC population (CD201^+^CD150^+^KL) and increased in fractions with lower HSC enrichment, in line with previous reports (Fig. S1B)(Dever et al., 2016). One week post editing (day 10 of culture), most cells in the expansion cultures remained c-Kit^+^ and Lin^-^, with a majority also expressing CD150 (Fig. 1D). Although the initial starting population of CD201^+^CD150^+^KL cells represented only 14.7% of expanded HSCs, absolute quantification revealed a 8.9-fold expansion (Fig. 1E).

To validate functional recovery of edited SCID HSCs, we transplanted expanded bulk HSC cultures into irradiated CB17/SCID recipients 7 days post-editing. B220^+^ B cells as well as CD4^+^ and CD8^+^ T cells could be detected in peripheral blood (PB) samples from 4 weeks post-SCT (Fig. 1F). Spleens of transplanted mice contained high fractions of B and T cells (B220^+^ 36%, CD4^+^ 16%. CD8^+^ 6% of splenocytes, Fig. S1C). We further found that thymocytes of transplanted mice were abundant with CD4^+^CD8^+^ double positive (DP), CD4^+^, and CD8^+^ single positive (SP) cells (Fig. S1D) and thymus histology showed cortical and medullary regions (Fig. S1E). The distributions of lymphocyte populations in the spleen and thymus were similar to those in age matched CB17/WT mice, suggesting orthotopic development of B and T lymphocytes. Secondary transplantations confirmed that LT-HSCs had been successfully edited in our gene correction model (Fig. 1F).

As mentioned above, the high frequency of indel and scid alleles in the transplanted HSPC population is a key limitation of this straightforward bulk expansion approach (Fig. 1C). To check how this distribution was reflected in mature cell lineages, we sequenced PB cells 16-20 weeks post-SCT. As expected, *Prkdc*^HDR^ frequencies were high in lymphocytes (B220^+^: 69%, CD4^+^: 70%, CD8^+^: 63%), suggesting at least monoallelic correction in these populations (Fig. 1G). Since noncorrected cells fail to complete lymphocyte development in this model, their high prevalence in the transplanted graft did not obstruct the rescue of these mature compartments. By contrast, myeloid cells, which are not subject to the same selective pressure, showed a high rate of on-target indels (*Prkdc*^indel^, 52%) and a low frequency of *Prkdc*^HDR^ alleles (14%, Fig. 1G). Off-target analysis of bulk-expanded HSCs showed a low but substantial prevalence of non-intended edits (Fig. 1H, Table S1).

While gene-edited HSCs effectively reversed the *Prkdc*^scid^ phenotype, these results indicate that most transplanted HSCs and their progeny contained unintended perturbations. The low allelic chimerism of *Prkdc*^HDR^ and high abundance of *Prkdc*^indel^ among myeloid cells demonstrate the challenge of ensuring that all hematopoietic cells are supplied by a genetically defined population of edited HSPCs. The potentially negative consequences of on-target indels has also recently been highlighted in other gene correction models (Wilkinson et al., 2021). This drawback inspired us to establish a single cell HSC expansion system that would allow sequence-based selection of *bona fide* HSCs at the clonal level.

### CD150^+^CD201^+^CD48^-^KSL cells contain clones with long term expansion potential

Single cell expansion of edited HSCs requires the identification of clones with prospective long-term (LT) expansion potential within a population of precultured cells. Cell surface markers are particularly useful since they permit flow cytometric profiling and simultaneous cloning via fluorescence-activated cell sorting. However, HSC marker expression undergoes a dynamic shift over the course of *ex vivo* expansion (Cheng and Lodish, 2005; Noda et al., 2008).

To address this issue, we leveraged index sorting analysis to identify HSC markers that predict LT expansion of HSC clones. Fresh CD34^-^CD150^+^c-Kit^+^Sca-1^+^Lin^-^(CD34^-^CD150^+^KSL) HSCs were cultured for 10 days, after which KSL cells were subjected to index sorting. We used C57BL/6-derived HSCs for these experiments, since this was the background of mice used to optimize our HSC culture system (Wilkinson et al., 2019) and is widely used in the field. Marker profiles of each sorted KSL cell were compared with HSC colony formation after 14 days. Expression of a total of six HSC markers within the KSL population, divided into two panels (CD34, CD48 and CD105; as well as CD135, CD150 and CD201), were evaluated (Fig. 2A). Colony formation was observed in 17.1% of sorted KSL clones (set 1: 16.2%, set 2: 19.1%), mainly from clones within the CD48^-^, CD150^+^ and CD201^+^ KSL populations (Fig. 2B). Quantification of expression levels confirmed significantly higher expression of CD150 and CD201 as well as lower expression of CD48 among colony-forming HSCs (Fig. 2C-D). CD135 expression was lower in colony-forming HSCs (Fig. 2C), however, the small absolute difference in expression precludes the use of this marker for effective gating.

**Figure 2.**
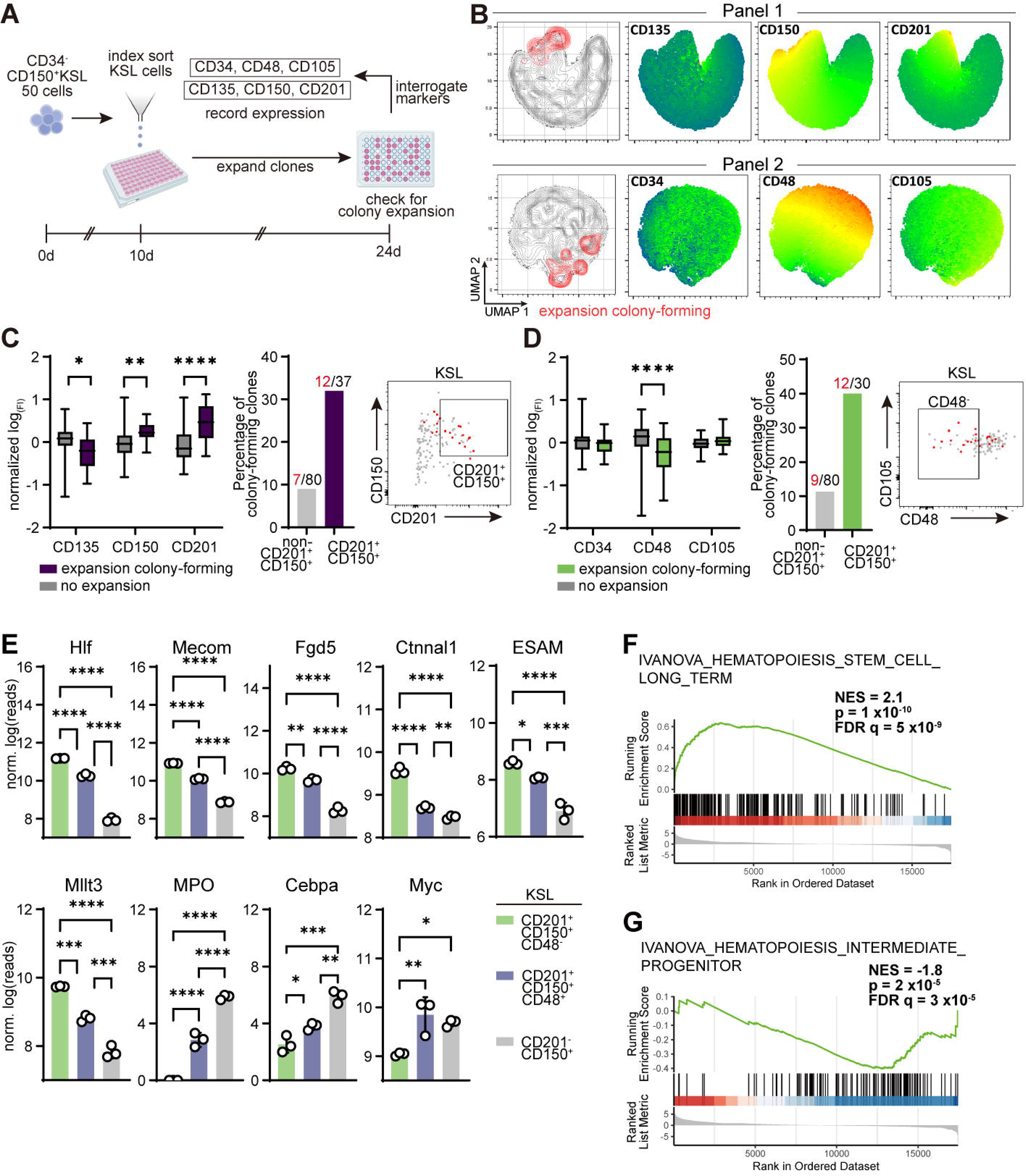
Identification of a surface marker combination for long-term (LT) expanding HSC clones. **(A)** Experimental setup. **(B)** Uniform Manifold Approximation and Projection (UMAP) representation of sorted KSL clones with overlay of panel 1 (upper) and 2 (lower) surface markers. Expansion colony-forming clones are indicated in red. **(C-D)** Quantification of markers associated with colony expansion. Left: Fluorescence intensity (FI) measured at index sorting. Data presented as log-transformed and normalized to mean. Box plots with whiskers showing minimum and maximum. Center: Fraction of clones of the indicated phenotype showing LT expansion. Right: Representative FACS sorting plots, LT-expanding clones indicated in red. **(C)** Panel 1 (n= 110 clones); **(D)** panel 2 (n=117 clones). Multiple Mann-Whitney tests with FDR correction. **(E)** RNAseq expression profiles of select HSC- and progenitor-associated genes. Error bars represent SD. One-way ANOVA with Tukey’s post-test. **(F-G),** Gene set enrichment analysis (GSEA) of differentially expressed genes in CD201^+^CD150^+^CD48^-^KSL **(F)** and CD201^-^CD150^+^KSL **(G)** cells. *P<0.05, **P<0.01, ***P<0.001, ****P<0.0001.

We next performed RNA sequencing to characterize the populations defined by these markers in 10-day bulk-expanded HSPCs (Fig. S2A). Comparing global expression profiles, we found the greatest difference between CD201^+^CD150^+^CD48^-^ and CD201^-^CD150^+^ KSL cells, with CD201^+^CD150^+^CD48^+^KSL cells representing an intermediary phenotype (Fig. S2B). This representation was mirrored in the expression profiles of canonical genes related to hematopoiesis: transcripts of HSC-associated genes, such as *Hlf*, *Mecom* and *Fgd5,* were more abundant in CD201^+^CD150^+^CD48^-^KSL cells, whereas downstream progenitor-associated genes (*MPO*, *Cebpa*) were upregulated in CD201^-^CD150^+^ KSL cells (Fig. 2E). Enrichment analysis confirmed that the transcriptional phenotype of CD201^+^CD150^+^CD48^-^KSL cells was similar to that of LT-HSCs (Fig. 2F), while CD201^-^CD150^+^KSL cells were similar to progenitor cells (Fig. 2G). GO term enrichment pointed to a proliferating state of CD201^-^CD150^+^KSL cells, with several enriched mitosis- and translation-related pathways (Fig. S2D), matching our previous observation that progenitor cells proliferate more rapidly than primitive HSCs in culture (Fig. 1E).

These results indicate that CD150^+^CD201^+^CD48^-^KSL cells possess LT expansion potential and retain a transcriptional phenotype associated with *bona fide* HSCs after extended culture. We thus considered these cells suitable for single cell cloning and expansion. However, we found that the CD201^+^CD150^+^CD48^-^ expression profile was lost in single clone-derived colonies generated from this population after 14 days, suggesting that repopulating activity had been compromised (Fig. S2E). Since we have shown previously that HSC marker expression is preserved in bulk cultures even after 28 days of expansion (Wilkinson et al., 2019), we reasoned that the single cell cloning step and expansion conditions, rather than the total length of *ex vivo* expansion, were not supported by our expansion system.

### Soluplus is a superior alternative to PVA for single cell HSC expansion

Having established that HSC activity among bulk cultured cells is enriched in the CD150^+^CD201^+^CD48^-^KSL population but that PVA-based culture conditions poorly supported their clonal expansion after re-sorting, we sought to improve clonal expansion culture conditions by screening alternative serum replacement compounds. We cultured 50 freshly isolated CD34^-^KSL cells in media supplemented with recombinant albumin, PVA and 7 different polymers and evaluated cell growth after one week. Of all compounds tested, only Soluplus led to comparable levels of proliferation as PVA and recombinant albumin (Fig. S3A). Soluplus is an amphiphilic polyvinyl caprolactam-acetate polyethylene glycol (PCL-PVAc-PEG) graft copolymer approved for clinical use as a drug solubilizer (Linn et al., 2012). To identify the most suitable concentration, we performed transplantations with HSCs grown in titrated concentrations of Soluplus. Sixteen-week chimerism was comparable in all dose groups (Fig. S3B). Supplementation with 0.2% Soluplus occasionally led to mild micelle formation in cultures, which obscured the visibility of cells. Based on our assay, we selected a concentration of 0.1% Soluplus for our expansion system.

To validate the cell growth supporting properties of Soluplus, we directly compared single HSC expansion conditions using Soluplus and PVA. Freshly isolated, single CD34^-^CD150^+^KSL cells from C57BL/6 mice were cultured in individual wells on 96-well plates. After 19 days, expansion was evaluated by flow cytometric profiling including cell viability and HSC marker expression (Fig. 3A). Cell viability was higher in clones cultured in Soluplus-supplemented medium, as measured by propidium iodide (PI) exclusion staining (Fig. S3C). Accordingly, the percentage of cloned HSCs forming viable cell colonies (i.e. >20% live cells) was higher under Soluplus expansion conditions (Fig. 3B). Furthermore, we found that Soluplus supplementation was associated with a higher retention of HSC marker expression. In particular, the fraction of CD201^+^CD150^+^KSL cells was higher in clones cultured in Soluplus-containing medium, suggesting that Soluplus was superior in expanding phenotypically primitive HSCs (Fig. 3C, S3D).

**Figure 3.**
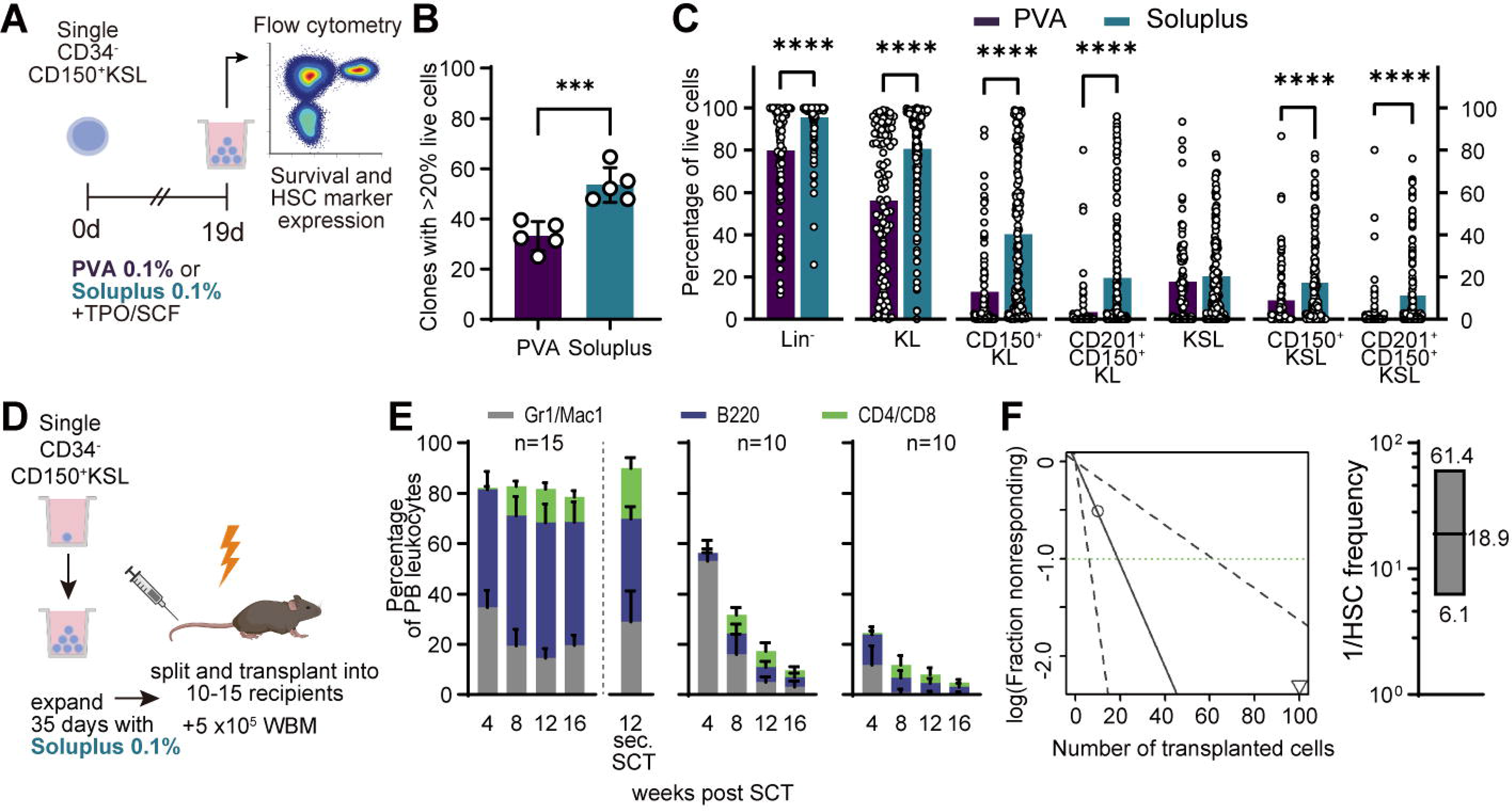
Optimization of polymer-based cultures for single cell HSC expansion. **(A)** Scheme of experimental setup. **(B)** Percentage of colonies with ≥ 0% live cells (n=5 experiments). Unpaired, two-tailed t-test. **(C)** Percentage of phenotypic HSC populations in live colonies cultured in PVA (n=94)- and Soluplus (n=155)-based media. Multiple Mann-Whitney tests with FDR correction. **(D)** Schematic of split clone transplantation. **(E)** Donor PB chimerism and lineage distribution in 3 recipient groups transplanted with split clones. Secondary SCT was performed with the group showing highest chimerism, data shown to the right of the dashed line. **(F)** Left: ELDA output of HSC frequency calculation. Right: Box plot represents calculated reciprocal mean, upper and lower limits of HSC frequency. Error bars represent SD. ***P<0.001, ****P<0.0001.

We next asked if HSC clones cultured in Soluplus medium produce functional HSC grafts *in vivo*. Freshly isolated CD45.1^+^CD34^-^ CD150^+^KSL HSCs were cloned and cultured for 35 days. Three clones containing 35%, 17% and 6% CD201^+^CD150^+^KSL cells were selected for split-clone transplantation into 10 to 15 CD45.2^+^ recipients against 5 x10^5^ WBM cells (Fig. 3D). All recipients showed multilineage LT engraftment (≥1% chimerism) in PB samples despite the high number of recipients per clone (Fig. 3E). Secondary transplantations from pooled bone marrow of highly chimeric mice showed successful engraftment of CD45.1^+^ cells in all secondary recipients (Fig. 3E).

To quantify the potential for single HSC expansion with Soluplus, we performed a limiting dilution assay (LDA) with a CD34^-^CD150^+^KSL clone expanded for 28 days (6.37 x10^5^ cells) and containing 84% of CD201^+^CD150^+^KSL cells (Fig. S3E). We observed multilineage chimerism of ≥ 1% in all dose groups including from just 10 cells (in 2/5 recipients, Fig. S3F-G). Based on these results, we estimated a mean HSC frequency of 1/18.9 (confidence interval (CI) 1/6.1-1/61.4) in the culture using extreme limiting dilation analysis (ELDA, Fig. 3F)(Hu and Smyth, 2009) and determined that the initial HSC had expanded >33,000-fold (range 10,375- to 104,426-fold corresponding to frequency CIs) under our culture conditions. Thus, our results suggest that Soluplus is superior to PVA in supporting efficient expansion of single HSCs. Based on these encouraging results, we attempted to expand precultured and gene-edited HSC clones.

### Soluplus enables single cell expansion of edited HSCs

To evaluate HSC gene editing and clonal expansion with single allele resolution, we developed a strategy that targets Protein Tyrosine Phosphatase Receptor Type C (*Ptprc*), a cell surface protein. Two alleles of the *Ptprc* gene are common among major inbred mouse strains: *Ptprc*^a^ and *Ptprc*^b^, which code for CD45.1 (Ly5.1) and CD45.2 (Ly5.2), respectively. The CD45.1 allele is expressed in SJL/J and STS/A strains, while C57BL/6 and BALB/c strains share the CD45.2 allele (Komuro et al., 1974). Sequence diversion between these two alleles amounts to 12 base differences that result in 5 amino acid substitutions (Zebedee et al., 1991). The epitope of CD45.1- and CD45.2-binding antibody clones A20 and 104 is defined by a single base difference at codon 302 (based on reference transcript NM_001111316.2, Fig. 4A)(Mercier et al., 2016). We leveraged the low complexity of this single nucleotide polymorphism (SNP) to simultaneously identify and clone gene-edited HSCs, followed by single cell expansion and transplantation (Fig. 4B). To this end, we knocked in the CD45.1-specific SNP variant (A→ G, p.K302E) into the *Ptprc* gene of CD45.2^+^ HSCs (Fig. S4A).

**Figure 4.**
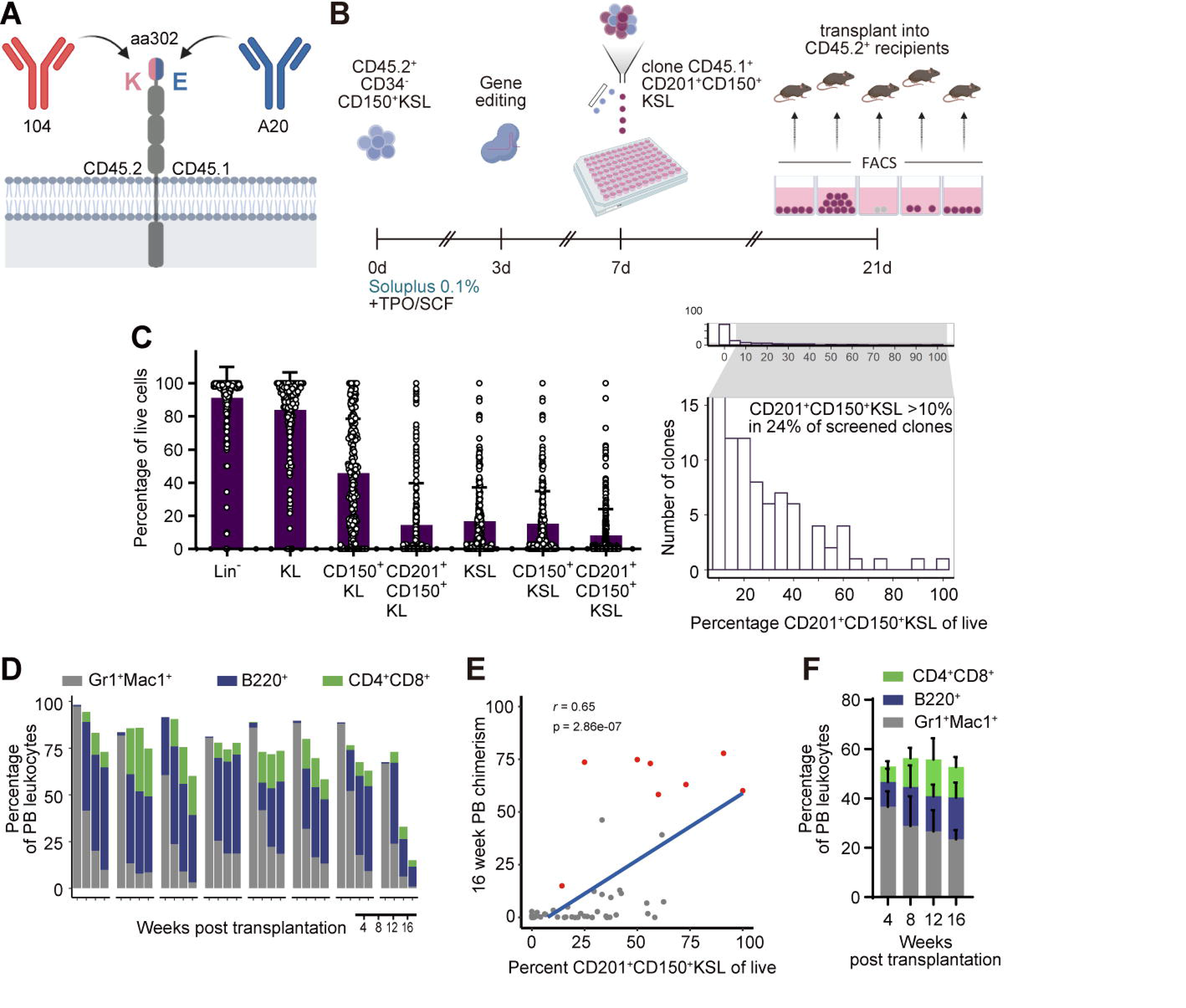
Single cell cloning of gene-edited functional HSCs. **(A)** Schematic showing the extracellular domain of CD45 with allele-specific antibody clones 104 and A20 and the epitope-defining amino acid. **(B)** Experimental setup of the single cell editing and expansion experiment. **(C)** Left: Fractions of CD201^+^CD150^+^KSL cells in single cell-derived cultures 14 days after cloning (n=261 clones). Right: Histogram of CD201^+^CD150^+^KSL cell frequency. Zoomed-in region shows clones with >10% CD201^+^CD150^+^KSL cells. **(D)** CD45.1^+^ donor PB chimerism and lineage distribution in single recipients with long-term (LT) engraftment 5% and ≥ multilineage reconstitution (n=8). **(E)** Linear correlation plots of CD201^+^CD150^+^KSL cell frequency and 16-week donor chimerism. Red dots indicate LT repopulating and multilineage clones. Pearson correlation. **(F)** CD45.1^+^ PB chimerism and lineage distribution in secondary recipients (n=5). Error bars represent SD.

CD34^-^CD150^+^KSL HSCs from CD45.2^+^ C57BL/6 mice were cultured in Soluplus-HSC expansion medium for 3 days, after which we targeted *Ptprc* for allele conversion. Four days after editing, 20% (±5.8%) of cells had converted to the CD45.1^+^CD45.2^-^ phenotype (Fig. S4B). As with the SCID model, conversion rates were lower in the primitive CD201^+^CD150^+^KSL fraction (Fig. S4C). We started 570 single cell cultures from the CD45.1^+^CD201^+^CD150^+^KSL population. 14 days later, cell proliferation could be observed in, and appropriate flow cytometric data could be obtained from, 46% of all sorted clones (261/570). Surface marker expression was heterogenous, with 24% (63/261) containing at least 10% of CD201^+^CD150^+^KSL cells (Fig. 4C). Fifty-one colonies were selected for transplantation into single CD45.2^+^ recipients. Donor chimerism of ≥5% was observed in 29/51 (57%) and 17/51 (33%) ≥ recipients 4 and 16 weeks after transplantation, respectively. Among the recipients showing LT chimerism, multilineage reconstitution (myeloid, B cell and T cell lineages ≥ % of donor hematopoiesis) was observed in 8 recipients (16% of recipients) (Fig. 4D), while the remaining mice showed biased donor hematopoiesis (Fig. S4D). Linear correlation analysis of pre-SCT marker expression and 16-week chimerism revealed several parameters associated with LT engraftment, the strongest of which was the fraction of CD201^+^CD150^+^KSL cells in the transplanted graft (Fig. 4E, Fig. S4E). Secondary transplantations were performed with whole bone marrow cells from a highly chimeric primary recipient. Analysis of bone marrow cells revealed high chimerism of 77% within the KSL population (Fig. S4F). Sixteen weeks after secondary transplantation, multilineage PB donor chimerism was observed in all secondary recipients (Fig. 4F).

Together, these results established that HSCs can be gene edited and clonally expanded while maintaining their self-renewal properties using our expansion system. Our experiments also confirm the expression of CD201 and CD150 on expanded clones as predictive of LT engraftment. This approach therefore provided the framework for probing single HSC clones for on- and off-target edits prior to transplantation.

### Single cell expansion of edited HSCs permits the assembly of a genetically defined HSC graft

To explore this approach, we adopted our single cell expansion platform to the SCID immunodeficiency model. Unlike CD45 allele conversion, *Prkdc* editing does not produce a detectable marker on cell surface proteins. Therefore, the generation of a high proportion of *Prkdc*^HDR^ alleles prior to cloning is desirable to increase the total yield of transplantable candidates. As it has been reported that inactive *Prkdc* may shunt the DDR towards the HDR pathway in cell lines, increasing knockin rates (Riesenberg et al., 2019), we initially compared HDR efficiencies in *Prkdc*-deficient (CB17/SCID) and -proficient (CB17/WT) HSCs using our CD45 allele conversion assay. We did not observe an increase in the CD45.1^+^ fractions, suggesting that HDR frequencies are unaltered in SCID HSCs (Fig. S5A). In contrast, the population of CD45^-/-^ knockout cells was significantly increased in SCID HSCs, suggesting higher prevalence of indels due to non-HR-based repair. Sequencing of alleles in this population revealed a relatively higher proportion of large indels (≥5bp, Fig. S5B) in SCID HSCs, indicative of alternative end joining pathways such as microhomology-mediated end joining (MMEJ), which has been reported to be de-repressed in *Prkdc*-deficient cells (McVey et al., 2008).

Analogous to our previous experiments, we expanded SCID HSCs in Soluplus-supplemented expansion medium and edited SCID HSCs to correct the *Prkdc*^scid^ mutation. After editing, CD201^+^CD150^+^CD48^-^ KL cells were cloned by flow cytometry and expanded for 14 days (Fig. 5A). Since expression of CD201 and CD150 was predictive of LT engraftment, we first screened for colonies containing a CD201^+^CD150^+^KL population of over 10% and then checked for the presence of the corrected allele (*Prkdc*^HDR^) and absence of off-target mutations. Candidate clones were then combined and administered to a SCID recipient (Fig. 5A). Phenotypic profiling data could be obtained from 19% (384) of sorted clones (Fig. 5B). Of these, 26% (99/384) contained a population of CD201^+^CD150^+^KL HSCs 10%, and sequencing of all intended loci could be achieved in most of ≥ these clones (96/384). We detected *Prkdc*^HDR^ in 57% (55/96) of genotyped clones, and all corrected clones were free of off-target mutations at predicted sites (Fig. 5C). As a result, an average of 18 HDR^+^Off-target^-^ colonies were selected for transplant per experiment. Due to this selection step, the combined allelic composition of the selected clones was dominated by *Prkdc*^HDR^ alleles (67%, Fig. 5D). This stands in contrast to our bulk-transplant approach, in which indel alleles were most abundant (Fig. 1C). In PB samples from transplanted SCB17/SCID mice, we detected B and T lymphocytes from week 4 through week 20 post-transplant, confirming LT engraftment (Fig. 5E). In contrast to our bulk-transplant experiments, the *Prkdc*^HDR^ allele was highly prevalent not only in lymphoid, but also in myeloid cells (>60%, Fig. 5F). Notably, *Prkdc*^indel^ frequency was low in all PB lineages.

**Figure 5.**
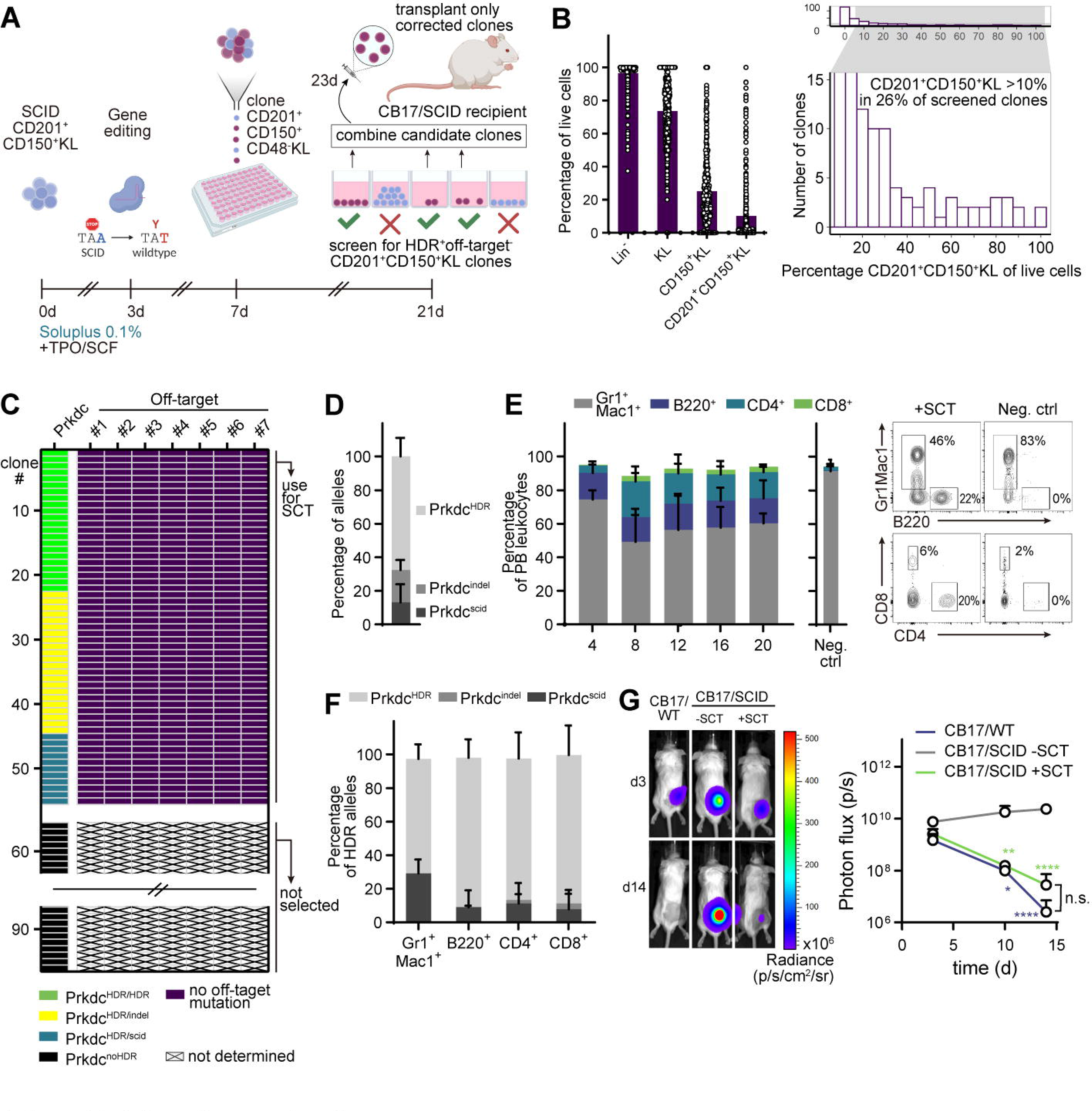
Autologous HSCT using single cell cloned gene-corrected HSCs is curative in a immunodeficiency mouse model. **(A)** Schematic of the single clone *Prkdc*^scid^ correction model. **(B)** Single cell SCID HSC expansion outcomes. Left: Frequencies of phenotypic HSC populations in screened colonies (n=384 from 3 experiments). Right: Histogram of CD201^+^CD150^+^KL cell frequency. Enlarged region shows clones with 10% CD201^+^CD150^+^KL cells. **(C)** Genotyping of candidate clones (n=96 clones, 3 ≥ experiments). Only clones with at least one HDR-corrected allele were sequenced at the off-target loci. **(D)** Allelic composition of the combined cell mixture at the edited *Prkdc* locus (n=3). **(E)** Frequencies of PB leukocytes in CB17/SCID recipients. Left: Lineage distributions in treated mice (n=3) and in recipients receiving only 2 x10^5^ CB17/SCID whole bone marrow cells (neg. ctrl., n=3). Right: representative FACS plots at 16 weeks post-SCT. **(F)** Allele frequencies in sorted PB cells 20 weeks post-SCT (n=3 mice from 3 experiments). **(G)** Xenograft transplantation assay. A549 cells expressing the luminescent reporter Akaluc were injected s.c. and tumor growth was tracked by *in vivo* imaging. Left: Representative images from CB17/WT, transplanted CB17/SCID, and untreated CB17/SCID mice 3 and 14 days after inoculation. Right: Quantification of luminescence over a 14-day period (CB17/WT: n=4, CB17/SCID-SCT: n=3, CB17/SCID+SCT: n=3). Two-way ANOVA and Tukey’s multiple comparison test. Error bars represent SD. *P<0.05, **P<0.01, ****P<0.0001.

Having achieved robust reconstitution of lymphoid cells, we asked if correction of the *Prkdc*^scid^ allele also led to development of a functional immune system. Double (CD4^+^CD8^+^) and single (CD4^+^ and CD8^+^) positive cells were detected among thymocytes of SCID recipients (Fig. S5C). Length diversity of the third complementary determining region (CDR3) in the T cell receptor gene is a direct function of *Prkdc* activity. We measured CDR3 region length distributions within multiple T cell receptor beta chain (*Tcrb*) gene families of splenic CD4^+^ T cells using spectratype analysis (Pannetier et al., 1993). Distribution profiles showed gaussian distribution patterns without oligoclonal spikes (Fig. S5D), suggesting that restored *Prkdc* activity permitted the generation of unbiased CDR3 regions. To confirm the status of immune cell function *in vivo*, we immunized transplanted CB17/SCID mice with a T-dependent antigen, nitroiodophenyl (NIP)-conjugated OVA (NIPOVA), intraperitoneally (i.p.)(Lee et al., 2019). NIP_30_-specific IgG and IgM levels were significantly elevated in the serum of transplanted CB17/SCID mice 19 days post-immunization (p.i.), indicative of a specific humoral immune response (Fig. S5E). On the other hand, untreated CB17/SCID mice showed no specific response. To assess cellular immunity, we inoculated mice with the human lung cancer cell line A549, hypothesizing that reconstitution of immunity would trigger xenograft rejection. Indeed, rejection of the injected cells was observed in transplanted mice only, whereas progressive tumor growth was detected in nontransplanted SCID mice (Fig. 5G). We therefore conclude that the molecular and functional hallmarks defining the *Prkdc*^scid^ phenotype had been reversed. In sum, these results illustrate that a functional graft can be assembled from individually expanded and profiled gene-edited HSC clones.

## Discussion

Here, we have introduced a fully defined culture system for clonal expansion and selection of gene-edited LT-HSCs. To our knowledge, this represents the first protocol to clonally expand, directly sequence, and transplant any functional primary adult tissue stem cells without the use of pluripotent intermediary cells. We believe our results not only have important implications for studying the genetics of hematopoiesis, but also highlight the potential of this approach for gene therapy. With the widespread interest in gene editing as a new therapeutic modality, concerns about hazardous on- and off-target mutations in edited cell products have become more visible (Sheridan, 2021). A string of recent reports have uncovered previously underappreciated lesions, such as kilo- and megabase-scale deletions as well as chromotrypsis, illustrating the potential risk of Cas9-based gene editing (Adikusuma et al., 2018; Boutin et al., 2021; Leibowitz et al., 2021). Our defined culture system addresses this concern in the murine model by enabling marker-free selection of edited LT-HSC clones with known on- and off-target mutational profiles.

Surface markers associated with HSC activity specifically in expanded HSCs have been identified in the human and murine systems, e.g. ITGA3 (Tomellini et al., 2019), EPCR (CD201) (Fares et al., 2017), and CD48 (Noda et al., 2008). We found that the CD201^+^CD150^+^CD48^-^KSL phenotype was most likely to contain long-term expanding clones, and that the addition of Soluplus facilitated the expansion of transplantable, *bona fide* HSC colonies to previous unattainable levels of over 30,000-fold. *In vivo*, this translated to high levels of chimerism in split-clone transplantations. The mechanism by which Soluplus, a biocompatible excipient for oral drug formulations, exerts its HSC-supportive properties remain undefined, yet it seems reasonable to assume that Soluplus enhances the solubility and stability of cytokines and other essential factors (Linn et al., 2012; Obata et al., 2014). The fact that many sorted clones did not produce colonies at all underlines the need for novel markers to further resolve LT-HSC activity in expansion cultures.

Previous studies have achieved molecular reversion of the *Prkdc*^scid^ mutation in Lin^-^ bone marrow cells, but low efficiencies have obstructed effective functional correction of immunodeficiency *in vivo* (Abdul-Razak et al., 2018; Rahman et al., 2015). This might be attributed to the higher occurrence of mutagenic repair events, such as MMEJ. Our demonstration that a sufficient corrected cell dose can be generated despite this challenging background highlights the utility of HSC expansion to propagate corrected cells *ex vivo*. Such an approach would also be highly applicable to settings where on-target mutations are detrimental and limit the curative potential of gene correction approaches, e.g. correction of the hemoglobin sickle allele (Wilkinson et al., 2021).

A possible limitation of single cell-expanded HSCs for SCT is the oligoclonal composition of the transplanted graft, which might raise concerns regarding clonal dominance. While these concerns are warranted, it is important to consider the impact of Cas9 gene editing on the clonal composition of HSC pools *in vivo*. Cas9-mediated DSBs induce activation of p53 in edited HSCs, restricting clonal diversity (Dever et al., 2016; Schiroli et al., 2019). A recent study by Ferrari, Jacob et al. has shown that the long-term repopulating graft arising from a Cas9-edited human HSC pool is dominated by less than 10 clones after transplantation into NSG mice (Ferrari et al., 2020). Similarly, Sharma, Dever et al. have reported that a median of 2 clones contribute to 50% of allele diversity in an *HBB* gene editing model (Sharma et al., 2021). Although these observations obtained from xenograft models may not accurately inform our understanding of clonal dynamics in the autologous setting, they do reflect the negative impact of Cas9 gene editing on HSC clonal diversity. Thus, one can speculate that our screening process based on the CD201^+^CD150^+^CD48^-^KSL phenotype selects for clones that would have dominated long-term hematopoiesis even if the bulk population had been transplanted.

In summary, we have developed an easily adoptable and powerful clonal expansion platform for precise genetic and functional interrogation of HSCs at the single cell level.

## Author contributions

Conceptualization, H.J.B. and S.Y.; Methodology, H.J.B., R.I., M.S.J.L., S.Y.; Investigation, H.J.B, R.I., M.S.J.L., S.Y.; Formal Analysis, H.J.B., R.I., M.S.J.L., A.W., D.K., T.K., C.C., A.T., Y.O.; Writing – Original Draft, H.J.B.; Writing – Review & Editing, A.W. and D.K.; Resources, A.T., Y.O.; Funding Acquisition, S.Y.; Supervision, S.Y.

## Supporting information

Key Reagents

## Acknowledgments

We thank the University of Tokyo Institute of Medical Science (IMSUT) FACS core laboratory for expert technical assistance. We thank T. Sano, Dr. K. Kolter and Dr. F. Guth of BASF for providing expert knowledge regarding the polymers used in this study, and K. Woltjen for fruitful discussions. We further thank Dr. M. Muratani and Tsukuba i-Laboratory LLP for sequencing services (https://www.tsukuba-i-lab.com). This work was supported by the German Research Foundation (BE 6847/1-1 to H.J.B.), the Japan Society for the Promotion of Science (JSPS; #20K16234 to M.S.J.L., #21F21108 and #20K21612 to S.Y.), the Kay Kendall Leukaemia Fund (A.C.W.), the Japan Science and Technology Agency (JST; #18071245 to C.C.), and the Japanese Agency for Medical Research and Development (AMED; #21bm0404077h0001 and #21bm0704055h0002 to S.Y.).

## MATERIALAND METHODS

### Resource availability

#### Lead contact

Further information and requests for resources and reagents should be directed to Satoshi Yamazaki.

#### Materials availability

This study did not generate new unique reagents.

#### Data and code availability

The RNAseq data and data belonging to figures have been uploaded to a Dryad repository and will be published upon acceptance of the manuscript (DOI: 10.5061/dryad.m905qfv2f).

### Experimental model and subject details

#### Mice

Male C.B-17/Icr-^+/+^Jcl wildtype (CB17/WT) and male C.B-17/Icr-^scid/scid^Jcl (CB17/SCID) mice were obtained from Clea Inc., Japan. C57BL/6NCrSlc (Ly 5.2, CD45.2) mice were purchased from SLC Inc., Japan. C57BL/6-Ly5.1 (Ly 5.1, CD45.1) mice were purchased from Sankyo Labo, Japan. All mice were obtained at age 8-10 weeks and housed in specific-pathogen-free (SPF) conditions at up to 5 mice per cage, with free access to standard rodent feed and kept under a 12h light/12h dark cycle. All animal protocols were approved by the Animal Care and Use Committee of the Institute of Medical Science, University of Tokyo.

#### Primary cell and cell line cultures

All cell culture operations were conducted under sterile hoods. Cells were kept in an incubator (Panasonic) at 37°C and a constant CO2 fraction of 5%. For gene editing experiments, HSCs were cultured in hypoxia (5% F_i_O2). Cell concentrations were determined on a Countess II cytometer (Thermo Fisher Scientific) after staining with Turk’s staining buffer (bone marrow cells) or trypan blue dead stain solution. Male HSCs were cultured in a Ham’s F12 medium (Wako) supplemented with 10mM HEPES (Thermo Fisher Scientific), recombinant cytokines murine TPO (100 ng/ml, Peprotech) and SCF (10 ng/ml, Peprotech), as well as insulin-transferrin-selenium (ITS, Thermo Fisher Scientific, 1:100 dilution) and 1% Penicillin-Streptomycin-L-Glutamine (PSG, Wako). Recombinant human albumin (Albumin Biosciences), polyvinyl alcohol (PVA, 84% hydrolyzed, Sigma), Kollidon 12 FP, Kollidon 17 PF, Kollidon 90 F, Poloxamer 188 Bio, Poloxamer 407 Geismar, povidone and Soluplus (all BASF) were added at a concentration of 0.1% v/v (except for Soluplus titration experiments). PVA and Soluplus-supplemented media are designated PVA-HSC and Soluplus-HSC expansion medium, respectively. Polymers were added from prepared stocks of 10% w/v in ddH_2_O. Recombinant cytokines, ITS, PSG and polymers were freshly added to base media before each application. Cells were cultured on human fibronectin-coated 24-well dishes (Corning, for gene editing bulk expansion cultures) or on untreated U-bottom 96-well plates (TPP, for cultures starting with 1-50 cells). Human male epithelial lung cancer cell line A549 (ATCC) was cultured in DMEM (Wako) supplemented with 10% FBS (Thermo Fisher) and 1% Penicillin-Streptomycin (Wako). Cells were passaged after reaching 70-80% confluency. Transduction procedure with Akaluc-expressing lentivirus is described in the method detail section below.

### Method details

#### Murine HSC isolation

Male 8-10 week-old mice were sacrificed by cervical dislocation after isoflurane anesthesia. Pelvic, femur and tibia bones were isolated and crushed, and the obtained cell solution was filtered through a 48 µm nylon mesh and whole bone marrow cells were counted. Positive selection of it^+^ cells was performed with anti-APC magnetic-activated cell sorting (MACS, Miltenyi Biotec) antibodies after staining cells with c-Kit-APC antibody for 30 minutes. Enriched it^+^ cells were incubated with anti-Lineage antibody cocktail (consisting of biotinylated Gr1[LY-6G/LY-6C], CD11b, CD4, CD8a, CD45R[B220], IL7-Rα, TER119) for 30 minutes. This was followed by staining with CD34-FITC, α CD201-PE (for CB17 strains) or Sca-1-PE (for C57BL/6 strains), c-Kit-APC, streptavidin-APC/eFluor 780 and CD150-PE/Cy7 antibodies for 90 minutes. CD201^+^CD150^+^c-Kit^+^Lin^-^ (CD201^+^CD150^+^KL) cells and CD34^-^CD150^+^c-Kit^+^Sca-1^+^Lin^-^ (CD34^-^CD150^+^KSL) cells from CB17 and C57BL/6 bone marrows, respectively, were sorted via fluorescence-activated cell sorting (FACS) on a Aria II cell sorter (BD) using a 100 μm nozzle and appropriate filters and settings. Propidium iodide (PI) was used to exclude dead cells. For bulk expansion of HSCs before gene editing, 5000 cells were sorted into 1 ml of HSC expansion medium per well. Medium changes were not performed until gene editing (day 3 of culture). For single cell expansion of freshly isolated HSCs, single HSCs were sorted into individual wells on a 96-well U-bottom plate (TPP) prefilled with 200 µl of culture medium. Culture medium was changed on day 7 post-sort, after which complete media changes were performed every 2-3 days.

#### CRISPR/Cas9 gene editing

Seventeen micrograms of recombinant S. pyogenes Cas9 (S.p. Cas9 Nuclease V3, IDT) were complexed with single guide RNA (sgRNA, synthesized at IDT) at a molar ratio of 1:2.5 (104 pmol Cas9:260 pmol sgRNA) for 10 minutes at 25°C to form ribonucleoprotein (RNP) complexes. Sequences of sgRNA targeting *Prkdc*^scid^ (Prkdc_gRNA1) and *Ptprc*^b^ (CD45.2, Ptprc_gRNA1) are listed in Table S2. Expanded HSCs were washed twice with PBS, pelleted, and resuspended in 20 μl electroporation buffer P3 (Lonza). Cells were gently added to the RNP duplex. For knockin experiments, 200 pmol of single-strand oligonucleotide (ssODN) templates (synthesized at IDT, Table S2) were added to the cell-RNP suspension. The suspension was transferred to a single 20 µl electroporation cuvette on a 16-well strip (P3 Primary Cell 96-well-Nucleofector Kit, Lonza). Electroporation was carried out on a 4D nucleofector device (Lonza) using programs DI-100 (CB17 HSCs) and EO-100 (C57BL/6 HSCs). Cells were immediately recovered in pre-warmed medium and gently split-transferred into 3 wells on a human fibronectin-coated 24-well plate (Corning) at 1 ml per well. A medium change was performed one day after nucleofection, and further medium changes were performed every 2-3 days.

#### Indel and HDR quantification in bulk-expanded cultures

To quantify indel and HDR rates from bulk cultured cells, genomic DNA (gDNA) was extracted using NucleoSpin Tissue XS columns (Macherey-Nagel). DNA concentration was measured on a Nanodrop spectrophotometer (Thermo Fisher). 1-10 ng of gDNA was used for polymerase chain reactions (PCR), formulated as 0.5 μ forward and reverse primers (Prkdc_inner_F, Prkdc_inner_R), 10 μ 2X buffer, and 0.5 U of Gflex *Thermococcus* DNA polymerase (Takara) in a 20 μl reaction. The PCR reaction setup was as follows: initial denaturation 94°C, 60s; followed by 35 cycles of denaturation 98°C, 10s; annealing 60°C, 15s; extension 68°C, 45s; and final extension 68°C, 45s. PCR products were separated on a 1.5% agarose gel via electrophoresis and fragments corresponding to the expected amplification target were cut and gel-purified using the Wizard SV gel and PCR clean-up system (Promega). Fourty nanograms of purified fragment was subjected to Sanger sequencing (FASMAC, Japan) using the forward primer (Prkdc_inner_F). For assessment of HDR rates in bulk cultures, we used the web-based tool Inference of CRISPR edits (ICE, Synthego, https://ice.synthego.com). Sequences from non-edited HSCs were provided as negative control samples. Potential off-target sites associated with the designed *Prkdc* gRNA were identified using COSMID (Cradick et al., 2014) with up to 3 mismatches in the absence of indels in the seed sequence and 2 mismatches in the presence of one insert or deletion. All targets showing a score <3 were amplified using the same cycling conditions outlined above. For bulk expansion cultures, only “inner” primer pairs were used for PCRs (see Table S2). Sequencing was performed with either forward or reverse PCR primers (specified with “SEQ” in Table S2), except off-target site #6, for which a dedicated sequencing primer was designed (OT_06_SEQ). On- and off target indel frequency was calculated with the Tracking of Indels by

Decomposition (TIDE) algorithm (Brinkman et al., 2014).

#### Analysis of bulk expanded cells

Cell counting operations were performed on a Countess II cytometer (Thermo Fisher Scientific). For flow cytometric studies of bulk expansion cultures, i.e. HSPCs cultured in 1 ml of expansion media, a 100 µl aliquot was removed from the culture well, washed in PBS, and stained with lineage antibodies (PB- and BV421-conjugated against Gr1[LY-6G/LY-6C], CD11b, CD4, CD8, CD45R[B220], TER119), CD34-FITC, CD201-PE, Sca-1-APC/Cy7, c-Kit-APC, CD150-PE/Cy7 antibodies for 45 minutes. After washing once with PBS, cells were analyzed on a FACSVerse or FACSAria II flow cytometer (BD).

#### Peripheral blood analyses

For chimerism and lineage analysis, peripheral blood was drawn from mice by retro-orbital sinus sampling under general anesthesia. Red blood cells (RBC) in a sample of 40 µl were lysed in 1 ml of Ammonium-Chloride-Potassium (ACK, 0.15 M NH4Cl, 0.01 M KHCO3, 0.1 mM Na2EDTA) buffer for 15 minutes at room temperature. RBC lysis was repeated 2 times. Lysed blood cells were stained with Gr1-PE, CD11b-PE, CD4-APC, CD45R[B220]-APC/eFluor 780, CD8-PE/Cy7 for SCID mouse samples and with Gr1-PE, CD11b-PE, CD4-APC, CD8a-APC, CD45R[B220]-APC/eFluor 780, CD45.1-PE/Cy7 and CD45.2-BV421 for C57BL/6 mice samples for 30 minutes at room temperature. Cells were resuspended in 200 µl PBS/PI before recording events on a FACSVerse (BD) analyzer using the appropriate filters and settings.

#### Fluorescence-activated single cell index sorting

CD34^-^CD150 KSL cells were isolated from C57BL/6 mice and cultured on a 96-well dish in PVA-HSC expansion medium at 50 cells per well. After 10 days, cells were stained with antibodies against KSL (biotinylated lineage-antibodies (same mixture as used in ‘Murine HSC isolation’), followed by c-Kit-APC/H7, Sca-1-BV605, and streptavidin-BV421). Cells were then divided into two sets, and each set was stained with an antibody panel (panel 1: CD34-FITC, CD48-APC, CD105-PE; panel 2: CD135-APC, CD201-PE, CD150-PE/Cy7). We cloned single KSL cells using the index sorting function on a FACSAria II (BD). Well location and expression data of the sorted clones were extracted using the IndexSort plugin for FlowJo (Freier, 2020). Dimensionality reduction was performed with the UMAP plugin for FlowJo (McInnes et al., 2018). Expression data of the sorted clones were log-transformed and normalized to mean.

#### RNAseq analysis of expanded HSCs

Expanded cells were washed in PBS once and stained with the identical panel specified in ‘Analysis of bulk expanded cells’, except for CD48-FITC, which was used instead of CD34-FITC. Over 5000 cells per population were sorted into 1.5 ml tubes and subsequently lysed in 600 µl Trizol LS reagent (Thermo Fisher Scientific). RNA purification, library preparation and next-generation sequencing was performed by Tsukuba i-Laboratory, LLC. Libraries were prepared using the SMARTer cDNA synthesis kit (Takara) and the high-output kit v2 (Illumina), followed by sequencing on a NextSeq 500 sequencer (Illumina) at 2x 36 paired end reads. Data normalization and comparative analyses were performed with the DESeq2 package in R (Love et al., 2014). Genes with an adjusted p <0.05 were considered differentially expressed. Enrichment analysis of differentially expressed genes was performed with the gene set enrichment analysis (GSEA) functions in the clusterProfiler package (Wu et al., 2021) using molecular signature database (MSigDB) gene ontology biological process (C5 GO:BP) as well as chemical and genetic perturbations (C2:CGP) gene set collections. Heatmaps were generated with the ComplexHeatmap package (Gu et al., 2016).

#### Marker profiling of single cell expanded HSC clones

To measure HSC marker expression in single cell-derived clonal cultures, 30 μ aliquots of cells were recovered from HSC colonies (cultured in 200 μ), transferred to a 96-well staining plate, washed in PBS, and stained with either PB/BV421- or FITC-conjugated linage antibodies (Gr1[LY-6G/LY-6C], CD11b, CD4, CD8, CD45R[B220], TER119), CD201-PE, Sca-1-APC/Cy7, c-Kit-APC and CD150-PE/Cy7 for 45 minutes at room temperature. After washing with PBS on-plate, cells were resuspended in 200 μl PBS/PI and examined on a FACSVerse analyzer (BD) using appropriate filters μ and settings. Acquisition time was set to 20 seconds to ensure enough cells remained for genomic DNA extraction, if necessary.

#### Genotyping of Prkdc-edited single cell HSC clones

To quantify HDR in single cell expanded clones of *Prkdc*-corrected HSCs (genotyping), cells left over from HSC marker profiling (see previous section) were subjected to gDNA extraction using NucleoSpin Tissue XS columns (Macherey-Nagel). gDNA was eluted in 18 μ of ddH_2_O. Only clones containing CD201^+^CD150^+^KL cells were selected for genotyping. For genotyping of the *Prkdc* locus, a nested PCR strategy was employed. The outer PCR formulation was 5 μ of gDNA, 0.5 μ forward and reverse outer primers (Prkdc_outer_F, Prkdc_outer_R) and 12.5 μ of Q5 2X master mix (containing Q5 DNA polymerase, dNTPs and Mg^2+^) (New England Biosciences) in a 25 μl reaction.

The PCR reaction setup was as follows: initial denaturation 98°C, 30s; followed by 35 cycles of denaturation 98°C, 10s; annealing 65°C, 15s; extension 72°C, 45s; and final extension 72°C, 120s.

The PCR product was diluted 1:20 for the inner PCR reaction. For this reaction, 1 μ of diluted PCR product was combined with 0.5 μ forward and reverse nested primers (Prkdc_inner_F, Prkdc_inner_R) and 25 μ of Q5 2X master mix in a 50 μ reaction. 700 bp PCR products were purified and sequenced as outlined above (‘Indel and HDR quantification in bulk-expanded cultures’). A semi-nested PCR strategy was employed for sequencing of off-target edits, first amplifying all sites in a multiplex PCR reaction using outer and inner primers, followed by a second reaction for individual targets using inner primers only. The multiplex PCR reaction contained primers specific to all off-target loci. The formulation was 5 μ of gDNA, 0.25 μ outer and inner primers, 25 μ 2X buffer, and 1.25 U of Gflex *Thermococcus* DNA polymerase (Takara) in a 50 μ reaction. The PCR reaction setup was as follows: initial denaturation 94°C, 60s; followed by 35 cycles of denaturation 98°C, 10s; annealing 60°C, 15s; extension 68°C, 120s; and final extension 68°C, 45s. The PCR product was diluted 1:20 and used for amplification of individual off-target sites. These reactions were M inner primers, 12.5 μl 2X buffer, and μ 0.625 U of Gflex *Thermococcus* DNA polymerase (Takara) in a 25 μl reaction. The PCR reaction setup was as follows: initial denaturation 94°C, 60s; followed by 35 cycles of denaturation 98°C, 10s; annealing 60°C, 15s; extension 68°C, 120s; and final extension 68°C, 45s. After agarose gel separation, appropriate PCR products were purified and sequenced using the inner reverse primers. Sequencing was performed with either forward or reverse inner PCR primers (specified with “SEQ” in Table S2), except off-target site #6, for which a dedicated sequencing primer was used (OT_06_SEQ). Sequence traces were aligned to reference sequences to check for mutations.

#### Stem cell transplantation (SCT)

Cells in *Prkdc*-edited bulk HSC cultures were washed, resuspended in 300 µl PBS and divided into 3 aliquots for transplantation into three recipients. For experiments comparing PVA and Soluplus expansion conditions, single cell-derived clones were split into several aliquots for SCT into multiple recipients as stated in the main text. *Ptprc*-edited single cell clones were transplanted into a one recipient. For single cell *Prkdc*-corrected clones, candidate clones were selected based on HSC marker and genotyping and combined to a single dose for transplantation into one CB17/SCID recipient. For SCT with *Prkdc*-corrected cells, 0.2 x10^6^ whole bone marrow (WBM) cells from 10 week old male CB17/SCID mice were added to the graft as support to ensure survival immediately after myeloablation. For non-edited and *Ptprc*-edited C57BL/6-derived HSCs, 0.2 x10^6^ WBM competitor cells from C57BL/6 CD45.1^+^/CD45.2^+^ F1 mice were added unless stated otherwise in the main text. CB17/SCID and C57BL/6 mice were lethally irradiated with 2.5 Gy and 9 Gy, respectively, immediately prior to transplantation. Cells were injected via tail vein injection. Secondary bone marrow transplantations were performed by extracting WBM cells from the primary recipient and transplanting 1 x10^6^ cells into lethally irradiated secondary recipients.

#### CDR3 spectratyping

The spectratyping protocol originally published by Pannetrier et al. was followed with modifications by Ahmed et al. (Ahmed et al., 2009; Pannetier et al., 1993). Splenocytes were recovered by crushing freshly excised spleens between two glass slides (Matsunami). CD4^+^ lymphocytes were enriched using CD4 magnetic-activated cell sorting (MACS) positive selection according to anufacturer’s intructions (Miltenyi Biotec) and lysed in 300 µl Trizol reagent (Thermo Fisher) per 10^6^ cells. RNA was purified using the Direct-zol RNA Microprep kit (Zymo) and eluted in 15 µl ddH_2_O. 150 ng of RNA was subjected for cDNA synthesis using Superscript IV reverse transcriptase (Thermo Fisher) according to manufacturer’s instructions. 2 µ l of cDNA was used per Vβ PCR reaction. The PCR reaction was formulated as: 2 µl of cDNA, 20 pmol TCR constant region (TCR-Cb) and V β gene-specific primer (1 µM final concentration) (see Table S2), 10 μ 2X buffer (containing deoxynucleoside triphosphates (dNTPs) and Mg^2+^), and 0.5 U of Gflex DNA polymerase (Takara) in a 20 μl reaction. The PCR μ reaction setup was as follows: initial denaturation 94°C, 120s; followed by 40 cycles of denaturation 98°C, 10s; annealing 62°C, 30s; extension 68°C, 90s; and final extension 68°C, 600s. 5 µ l of PCR product was then used in a runoff reaction including a FAM-labeled TCR-Cb primer. The reaction mix was formulated as 5 µ l PCR product, 4 pmol 5’-FAM-labeled TCR constant region primer (TCR-Cb-FAM, 0.2 µ M final concentration), 10 μl 2X buffer (containing deoxynucleoside triphosphates (dNTPs) and Mg^2+^), and 0.5 U of Gflex *Thermococcus* DNA polymerase (Takara) in a 20 μ reaction. The PCR reaction setup was as follows: initial denaturation 94°C, 120s; followed by 20 cycles of denaturation 98°C, 10s; annealing 62°C, 30s; extension 68°C, 90s; and final extension 68°C, 300s. Ten microliters of the reaction mix were used for fragment sizing (performed at FASMAC, Japan). Fragment size analysis was performed on the Thermo Fisher Connect platform (https://apps.thermofisher.com/) using the peak scanner application. Relevant peaks were filtered and imported into Prism software (Graphpad) for further analysis and visualization (Miqueu et al., 2007). For Kolmogorov-Smirnov normality tests, a threshold level of 0.05 was selected to reject the hypothesis that data was normally distributed.

#### Immunization and ELISA assays

Mice at 20 weeks after SCT with *Prkdc*-corrected HSCs were immunized with 100 µg of nitroiodophenyl (NIP)-conjugated OVA (NIPOVA) (Biosearch Technologies) mixed 1:1 with aluminium hydroxide (Invivogen) intraperitoneally (i.p.). Blood samples were collected after 12 and 19 days post-immunization via retro-orbital sinus sampling. Serum was recovered by centrifugation of whole blood for 10 minutes at 5000g and stored at -20°C. For serum antibody detection, high binding 96-well microplates (Thermo Fisher) were pre-coated with NIP_30_-BSA (Biosearch Technologies) at 2 µg/ml concentration overnight. After blocking wells with 1% BSA/PBS solution, 1:5000 dilutions of serum samples were applied to the wells and incubated overnight at 4°C. Wells were washed with 0.05% PBS/Tween-20 (PBS-T) followed by incubation with horse radish peroxidase (HRP)-conjugated secondary antibodies against murine IgG and IgM (1:5000 dilution, Southern Biotech) for 2 hours. Enzymatic reaction was initiated by adding 100 µl of 3,3’,5,5’-Tetramethyl -benzidine (TMB) substrate solution (TCI) to each well, followed by termination of the reaction with 100 µl hydrochloric acid 1M (HCl, TCI). Absorbance readings were obtained on a microplate reader at 450 nm (Molecular Devices).

#### Xenograft transplantation assay

Human A549 cells were modified to constitutively express Akaluc, a firefly luciferase derivative with improved bioluminescent activity (Iwano et al., 2018). Cultured cells were transduced with a VSV-G pseudotyped lentiviral vector carrying an Akaluc-P2A-mNeonGreen transgene under the control of the human ubiquitin C (UbC) promoter at an MOI of 10. After 14 days, stably transduced cells were selected by sorting mNeonGreen-positive cells on a FACSAria II (BD). Xenograft transplantations were performed by subcutaneously injecting 5 x10^6^ cells in 100 µl of PBS into the flanks of recipient mice. Prior to intravital imaging, the fur above of the injection site was removed with household depilatory cream. After anesthesia, 50 µ l of Akalumine-HCl substrate (15 mM, Wako) were injected intraperitoneally and mice were placed in an IVIS in vivo imaging system (PerkinElmer). Images were acquired after 10-15 minutes using appropriate binning (1) and exposure settings.

### Quantification and statistical analysis

Details regarding employed statistical tests as well as number of subjects and groups are stated in the figure legends. Student’s t-tests, one- and two-way analysis of variance (ANOVA) were performed in Prism (version 9.1, Graphpad). Error bars denote standard deviations, unless otherwise stated in the figure legends. Statistical evaluation surrounding RNASeq analysis, correlation calculations between chimerism and CD201^+^CD150^+^KSL marker expression, as well as select visualizations of peripheral blood lineage distributions were performed in R version 4 (R Core Team, 2020) with the appropriate packages (outlined in key resources table). Pictograms and illustrations were generated with BioRender (https://www.biorender.com/).

## SUPPLEMENTAL FIGURE TITLES AND LEGENDS

**Figure S1.**
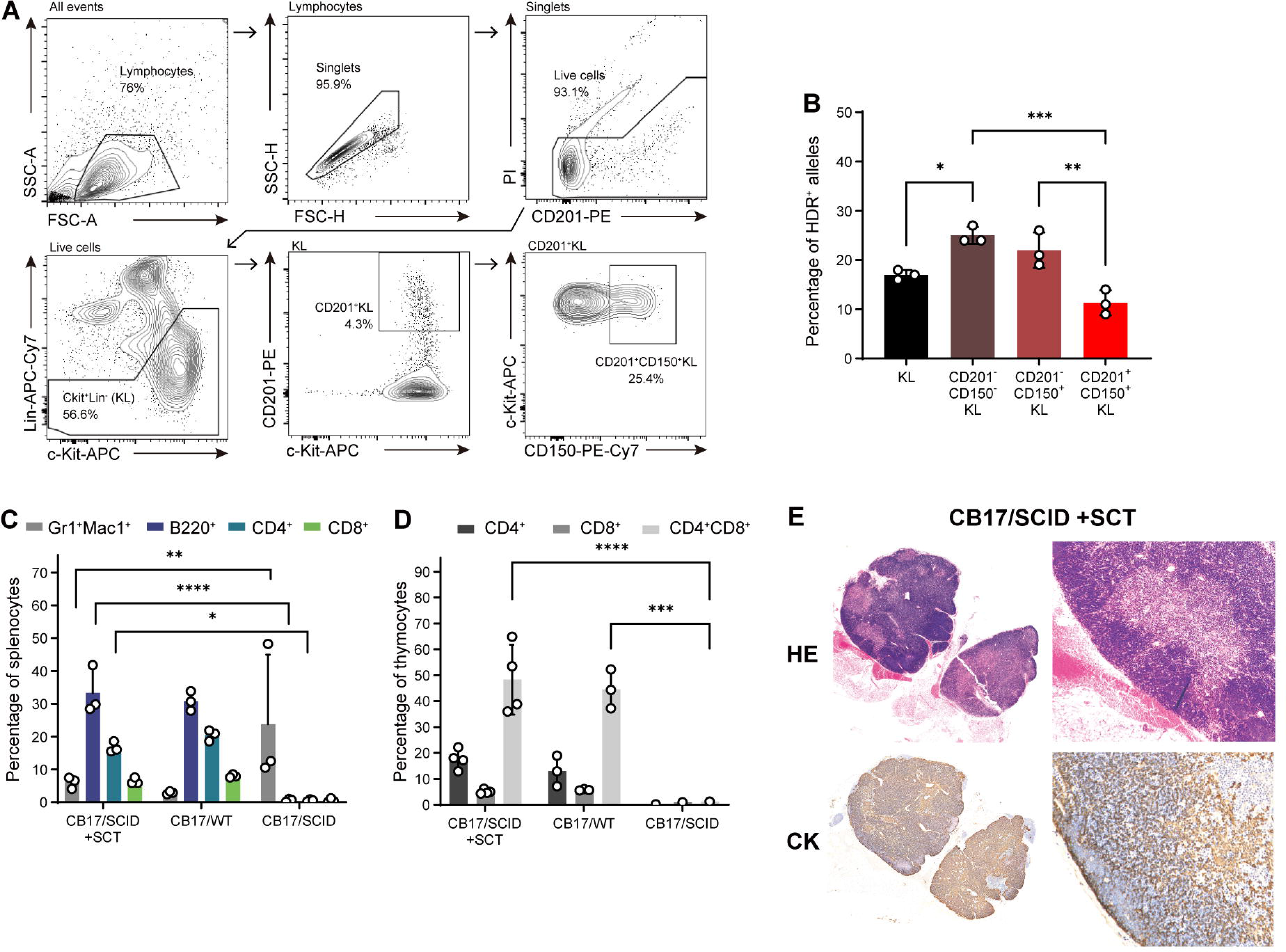
Functional correction of *Prkdc*^scid^ HSCs. Related to Fig. 1. **(A)** Gating strategy for the isolation of CD201^+^CD150^+^KL cells from CB17/SCID mouse BM. **(B)** Frequency of HDR^+^ alleles within phenotypically defined HSPC populations 7 days post gene editing (n=3 cultures). **(C)** Immunophenotype of splenocytes 20 weeks post-SCT (n=3 mice per group). **(D)** Frequencies of double (CD4^+^CD8^+^) and single positive (CD4^+^ and CD8^+^) thymocytes 20 weeks post-SCT. Data points represent individual mice. **(E)** Sections of thymi isolated from a CB17/SCID recipient transplanted with gene edited HSCs 20 weeks post-SCT. Upper panel: hematoxylin-eosin (HE), lower panel: cytokeratin (CK) stains. One-(B) and two-way(C, D) ANOVA with Tukey’s multiple comparison test. Error bars represent SD. *P<0.05, **P<0.01, ***P<0.001, ****P<0.0001.

**Figure S2.**
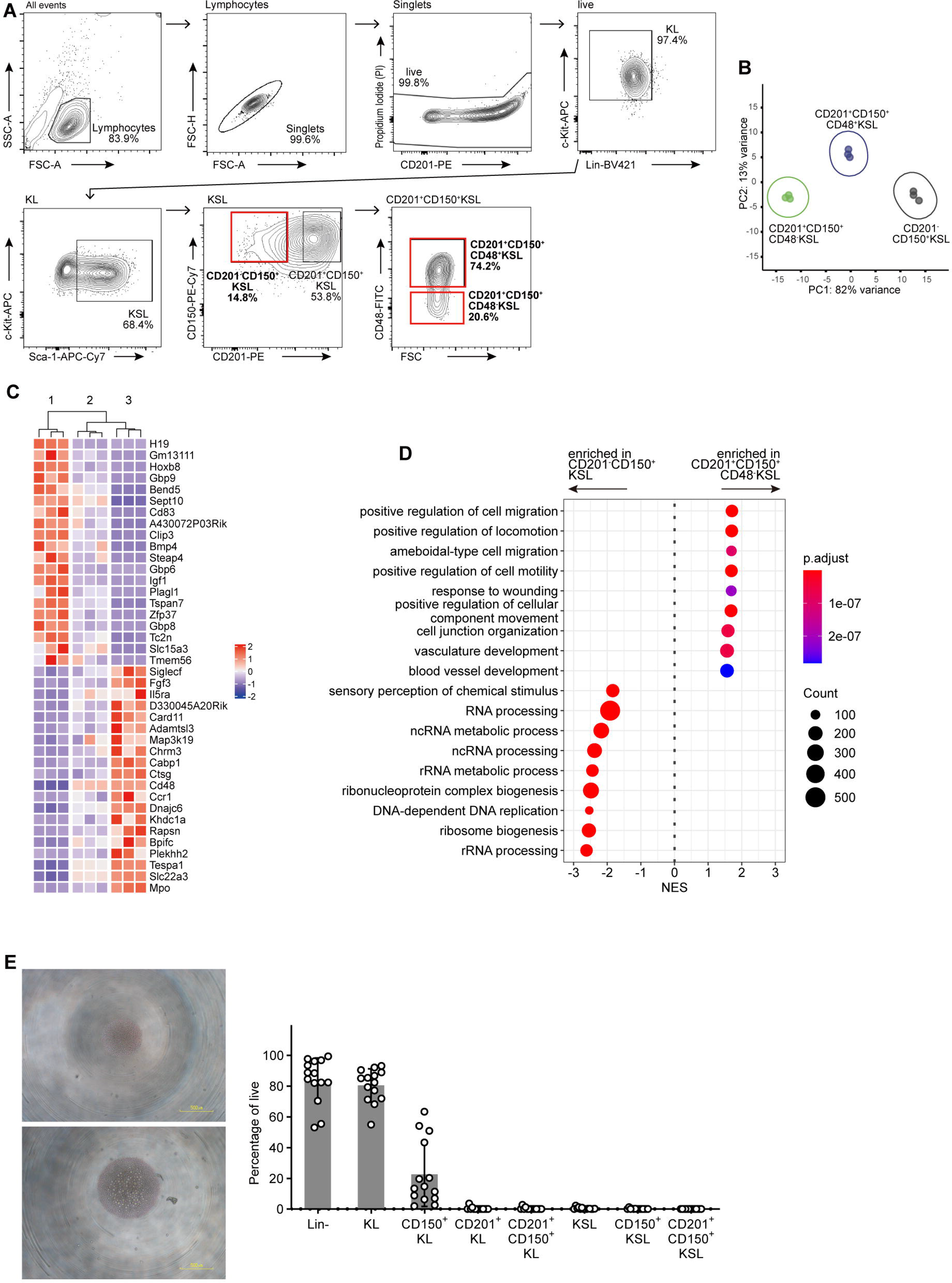
Transcriptional and immune phenotype of cultured, gene-edited HSCs. **Related to** Fig. 2**. (A)** Gating strategy applied for the isolation of CD201^+^CD150^+^CD48^-^KSL, CD201^+^CD150^+^CD48^+^KSL and CD201^-^CD150^+^KSL cells for RNAseq analysis on day 10 of bulk expansion. **(B)** Principal component analysis (PCA) of CD201^+^CD150^+^CD48^-^KSL, CD201^+^CD150^+^CD48^+^KSL and CD201^-^CD150^+^KSL cells (n=3 replicates). (**C)** Heatmap showing the top 40 differentially regulated (up- and down-regulated) genes in CD201^+^CD150^+^CD48^-^KSL (1) and CD201^-^CD150^+^KSL (3) populations, with expression data of CD201^+^CD150^+^CD48^+^KSL (2) cells (n=3 replicates). **(D)** Nine most significantly enriched GO terms (biological process) in CD201^+^CD150^+^CD48^-^ and CD201^-^CD150^+^KSL populations. **(E)** Expansion of precultured and cloned CD201^+^CD150^+^CD48^-^KSL cells. CD34^-^CD150^+^KSL cells were isolated and expanded in bulk for 10 days before cloning. The colonies generated from these clones were analyzed 14 days post-sort. Left: Representative images of single clone-derived HSC colonies 13 days post-sort. Right: Phenotypic fractions within expanded colonies, as a percentage of live cells (n=14).

**Figure S3.**
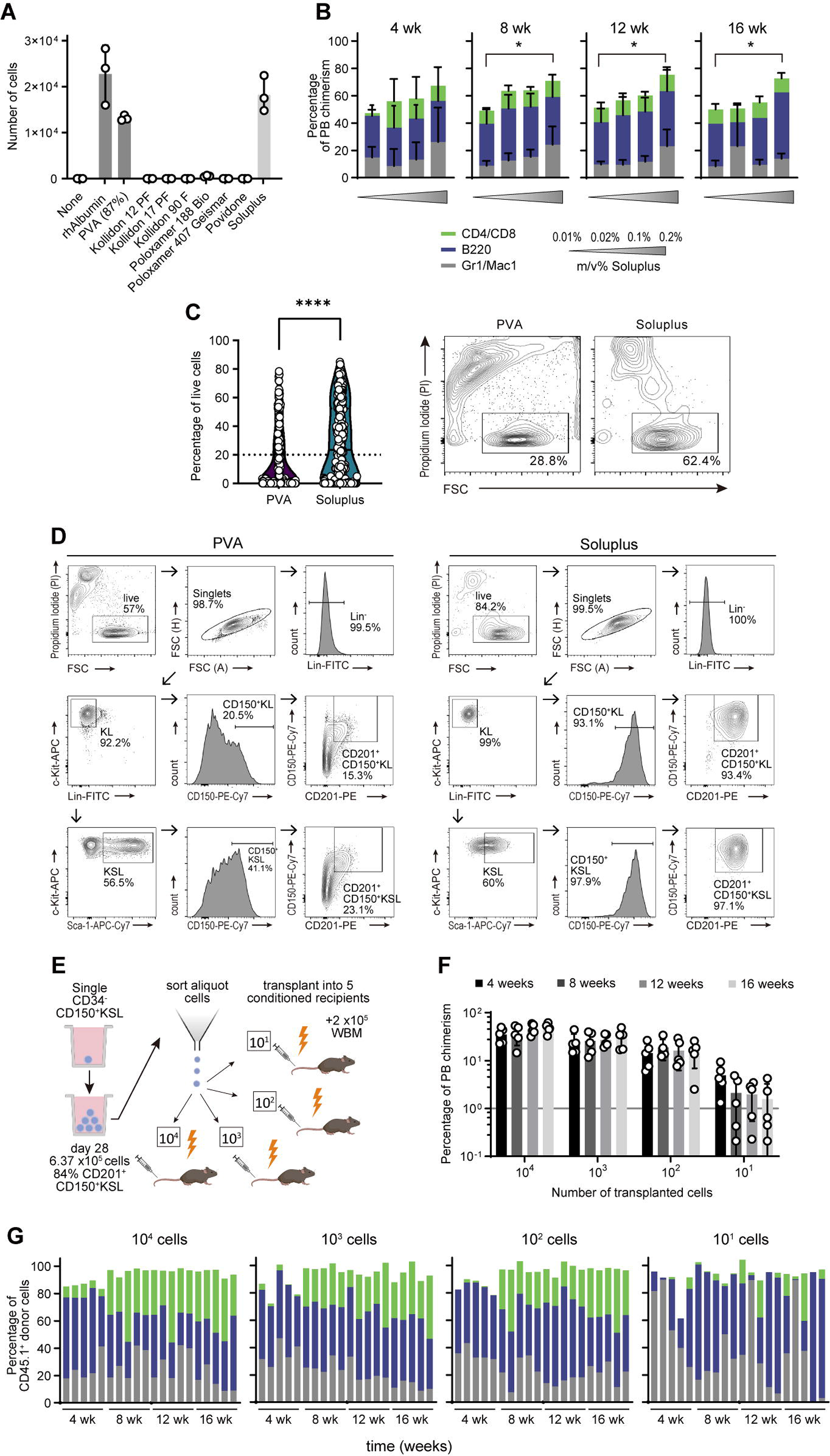
Titration of Soluplus supplementation and immune phenotype of Soluplus-expanded HSC clones. **Related to** Fig. 3**. (A)** Albumin replacement polymer screening for *ex vivo* expansion of murine HSCs. 50 freshly isolated C57BL/6 CD34^-^KSL cells were cultured in media supplemented with the indicated polymers for 7 days. Recombinant human albumin and PVA (87% hydrolyzed) served as positive controls. Concentration of all polymers was 0.1% (m/v). Total cells were counted to assess growth support (n=3 cultures). **(B)** Soluplus supplementation titration assay. Fifty C57BL/6-Ly5.1 (CD45.1^+^) CD34^-^CD150^+^KSL cells grown in titrated concentrations of Soluplus (0.01%, 0.02%, 0.1% and 0.2%) were expanded for 14 days and split-transplanted into CD45.2^+^ recipients (n=4 to 9 per group) against 2 x10^5^ CD45.1^+^/CD45.2^+^ whole bone marrow competitor cells. Peripheral blood (PB) chimerism and lineage distribution is shown. Supplementation with 0.2% Soluplus produced mild micelle formation in cultures, which was not toxic, but occasionally obstructed the visibility of cells. Two-way ANOVA with Tukey’s multiple comparison test. **(C)** Left panel: Percentage of viable cells in HSC colonies grown from freshly isolated single CD34^-^CD150^+^KSL cells after 19 days of culture in PVA (n=288)- and Soluplus (n=290)-based media, as evaluated by flow cytometry (%PI^-^ of all events). Right panel: Representative FACS plots of single-cell derived colonies (day 19). Two-tailed Mann-Whitney test. **(D)** Representative FACS plots of individual clones expanded in PVA- and Soluplus-supplemented media. **(E)** Schematic of limiting dilution assay (LDA). **(F)** Donor PB chimerism four to 16 weeks post-SCT (5 mice per group). Cutoff level (1%) denoted with gray line. **(G)** PB lineage distribution of CD45.1^+^ donor cells. Each bar represents an individual recipient. Error bars represent SD. *P<0.05, ****P<0.0001.

**Figure S4.**
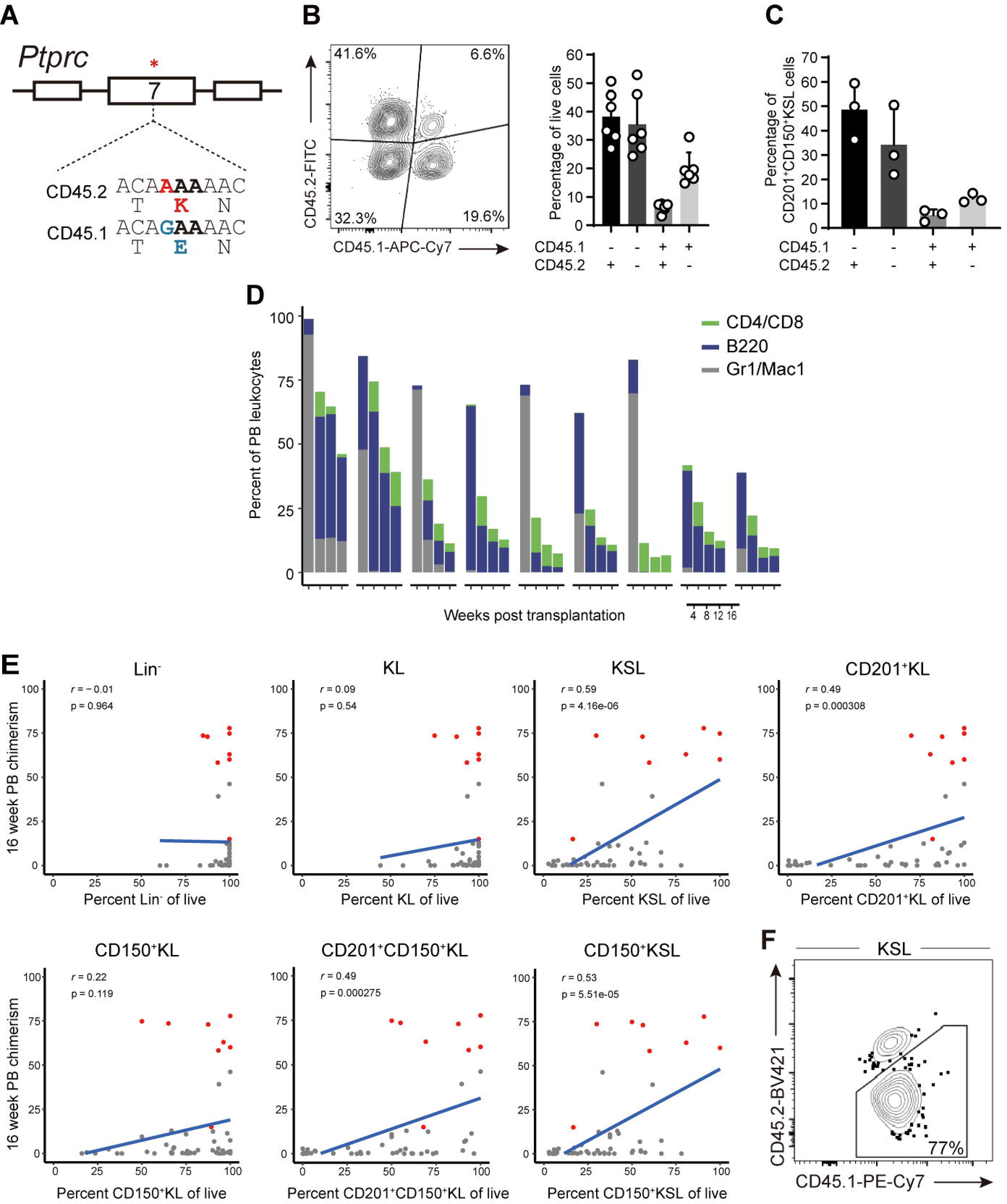
*Ptprc* allele conversion efficiency and single clone transplantation assays. Related to Fig. 4. **(A)** Genomic context of the *Ptprc*^a^ and *Ptprc*^b^ alleles. White boxes: exons. *denotes site of SNP. **(B)** Distribution of CD45 phenotypes among live cells 4 days post-editing. Representative FACS plot (left) and summary data (right, n=6 cultures). **(C)** Distribution of CD45 phenotypes among CD201^+^CD150^+^KSL cells 4 days post-editing (n=3 cultures). **(D)** CD45.1^+^ chimerism and lineage distribution in single recipients that did show LT chimerism 5% but did not display multilineage ≥ distribution (defined as each lineage ≥5%). Four to 16 weeks post-SCT. Related to Fig. 4d. **(E)** Correlation plots of different expansion culture phenotypes versus LT donor chimerism (16 weeks). Red dots indicate multilineage and LT repopulating clones. Pearson correlation. **(F)** Bone marrow CD45.1^+^ donor chimerism within the KSL compartment of a representative primary recipient. Error bars represent SD.

**Figure S5.**
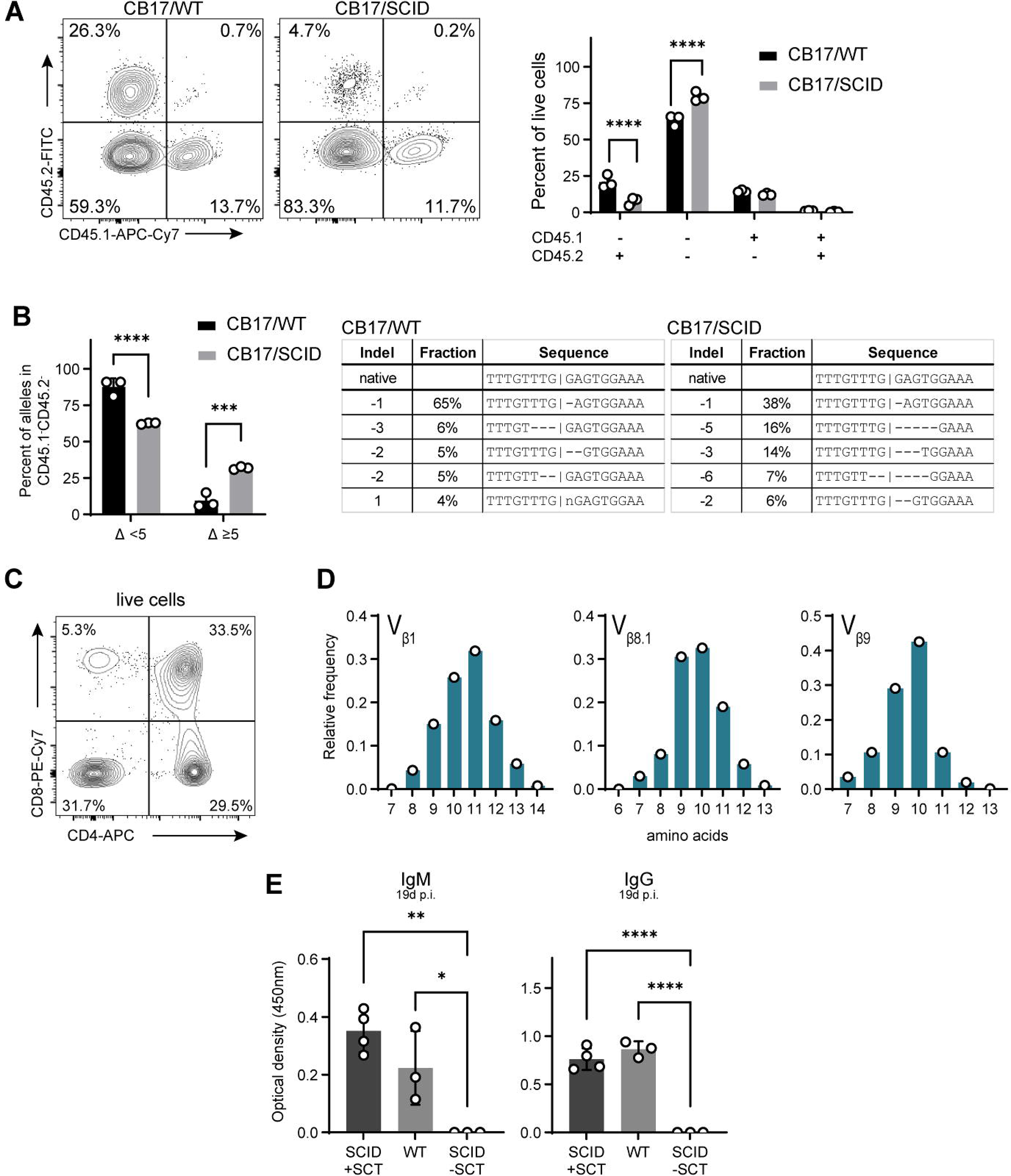
Functional rescue of SCID immunodeficiency with edited and single- cell expanded HSC graft. **Related to** Fig. 5**. (A)** HDR pathway efficiency in *Prkdc*-deficient HSCs. CB17/SCID and wildtype (CB17/WT) CD201^+^CD150^+^KL cells were isolated and cultured for 3 days, after which the CD45 allele conversion assay was performed (analogous to the experiment presented in Fig. 4A). Left: representative FACS plot showing the distribution of CD45 phenotypes among live cells HSCs 4 days post-editing. Representative FACS plot (left) and summary data (right, n=3 cultures). **(B)** Indel sizes at the *Prkdc* locus in the CD45.1^-^ CD45.2^-^ population of *Prkdc* WT and SCID HSCs 4 days post-editing. Left: Summary data of small (<5bp) and large (5bp) deletion events (n=3 cultures). Right: Representative≥ examples of indel distributions showing the most frequent alleles in both groups. **(C)** Representative FACS plot showing frequencies of double (CD4^+^CD8^+^) and single positive (CD4^+^ and CD8^+^) cells from the thymus of a CB17/SCID recipient 20 weeks after transplantation. **(D)** CDR3 length spectratype analysis of the Tcrb-V1, -V8.1 and -V9 genes in splenic CD4^+^ cells. Each bar represents the relative frequency of a CDR3 length species (n=1 mouse). **(E)** Serum levels of NIP_30_-specific IgG and IgM 19 days after immunization (CB17/SCID+SCT: n=4 mice; CB17/WT and treatment-naïve CB17/SCID: n=3). One-(E) and two-way (A, B) ANOVA with Tukey’s multiple comparison test. Error bars represent SD. *P<0.05, **P<0.01, ****P<0.0001.

## REFERENCES

Abdul-Razak, H.H., Rocca, C.J., Howe, S.J., Alonso-Ferrero, M.E., Wang, J., Gabriel, R., Bartholomae, C.C., Gan, C.H. V., Garín, M.I., Roberts, A., et al. (2018). Molecular Evidence of Genome Editing in a Mouse Model of Immunodeficiency. Sci. Rep. 8, 8214.

Adikusuma, F., Piltz, S., Corbett, M.A., Turvey, M., McColl, S.R., Helbig, K.J., Beard, M.R., Hughes, J., Pomerantz, R.T., and Thomas, P.Q. (2018). Large deletions induced by Cas9 cleavage. Nature 560, E8–E9.

Ahmed, M., Lanzer, K.G., Yager, E.J., Adams, P.S., Johnson, L.L., and Blackman, M.A. (2009). Clonal Expansions and Loss of Receptor Diversity in the Naive CD8 T Cell Repertoire of Aged Mice. J. Immunol. 182, 784–792.

Araki, R., Fujimori, A., Hamatani, K., Mita, K., Saito, T., Mori, M., Fukumura, R., Morimyo, M., Muto, M., Itoh, M., et al. (1997). Nonsense mutation at Tyr-4046 in the DNA-dependent protein kinase catalytic subunit of severe combined immune deficiency mice. Proc. Natl. Acad. Sci. U. S. A. 94, 2438–2443.

Boutin, J., Rosier, J., Cappellen, D., Prat, F., Toutain, J., Pennamen, P., Bouron, J., Rooryck, C., Merlio, J.P., Lamrissi-Garcia, I., et al. (2021). CRISPR-Cas9 globin editing can induce megabase-scale copy-neutral losses of heterozygosity in hematopoietic cells. Nat. Commun. 12, 4922.

Brinkman, E.K., Chen, T., Amendola, M., and Van Steensel, B. (2014). Easy quantitative assessment of genome editing by sequence trace decomposition. Nucleic Acids Res. 42, 1–8.

Cheng, C.Z., and Lodish, H.F. (2005). Murine hematopoietic stem cells change their surface phenotype during ex vivo expansion. Blood 105, 4314–4320.

Cradick, T.J., Qiu, P., Lee, C.M., Fine, E.J., and Bao, G. (2014). COSMID: A web-based tool for identifying and validating CRISPR/Cas off-target sites. Mol. Ther. - Nucleic Acids 3, e214.

Dever, D.P., Bak, R.O., Reinisch, A., Camarena, J., Washington, G., Nicolas, C.E., Pavel-Dinu, M., Saxena, N., Wilkens, A.B., Mantri, S., et al. (2016). CRISPR/Cas9 β-globin gene targeting in human haematopoietic stem cells. Nature 539, 384–389.

Fares, I., Chagraoui, J., Lehnertz, B., MacRae, T., Mayotte, N., Tomellini, E., Aubert, L., Roux, P.P., and Sauvageau, G. (2017). EPCR expression marks UM171-expanded CD34+ cord blood stem cells. Blood 129, 3344–3351.

Ferrari, S., Jacob, A., Beretta, S., Unali, G., Albano, L., Vavassori, V., Cittaro, D., Lazarevic, D., Brombin, C., Cugnata, F., et al. (2020). Efficient gene editing of human long-term hematopoietic stem cells validated by clonal tracking. Nat. Biotechnol. 38, 1298–1308.

Freier, C. (2020). indexSort: Retrieve Index Sorted Data and Visually Explore .fcs Date Files.

Genovese, P., Schiroli, G., Escobar, G., Tomaso, T. Di, Firrito, C., Calabria, A., Moi, D., Mazzieri, R., Bonini, C., Holmes, M.C., et al. (2014). Targeted genome editing in human repopulating haematopoietic stem cells. Nature 510, 235–240.

Gu, Z., Eils, R., and Schlesner, M. (2016). Complex heatmaps reveal patterns and correlations in multidimensional genomic data. Bioinformatics 32, 2847–2849.

Hu, Y., and Smyth, G.K. (2009). ELDA: Extreme limiting dilution analysis for comparing depleted and enriched populations in stem cell and other assays. J. Immunol. Methods 347, 70–78.

Iwano, S., Sugiyama, M., Hama, H., Watakabe, A., Hasegawa, N., Kuchimaru, T., Tanaka, K.Z., Takahashi, M., Ishida, Y., Hata, J., et al. (2018). Single-cell bioluminescence imaging of deep tissue in freely moving animals. Science 359, 935–939.

Komuro, K., Itakura, K., Boyse, E.A., and John, M. (1974). Ly-5: A new T-lymphocyte antigen system. Immunogenetics 1, 452–456.

Lattanzi, A., Camarena, J., Lahiri, P., Segal, H., Srifa, W., Vakulskas, C.A., Frock, R.L., Kenrick, J., Lee, C., Talbott, N., et al. (2021). Development of β-globin gene correction in human hematopoietic stem cells as a potential durable treatment for sickle cell disease. Sci. Transl. Med. 13.

Lee, M.S.J., Natsume-Kitatani, Y., Temizoz, B., Fujita, Y., Konishi, A., Matsuda, K., Igari, Y., Tsukui, T., Kobiyama, K., Kuroda, E., et al. (2019). B cell-intrinsic MyD88 signaling controls IFN- -mediated γ early IgG2c class switching in mice in response to a particulate adjuvant. Eur. J. Immunol. 49, 1433–1440.

Leibowitz, M.L., Papathanasiou, S., Doerfler, P.A., Blaine, L.J., Sun, L., Yao, Y., Zhang, C.Z., Weiss, M.J., and Pellman, D. (2021). Chromothripsis as an on-target consequence of CRISPR–Cas9 genome editing. Nat. Genet. 53, 895–905.

Linn, M., Collnot, E.M., Djuric, D., Hempel, K., Fabian, E., Kolter, K., and Lehr, C.M. (2012). Soluplus® as an effective absorption enhancer of poorly soluble drugs in vitro and in vivo. Eur. J. Pharm. Sci. 45, 336–343.

Love, M.I., Huber, W., and Anders, S. (2014). Moderated estimation of fold change and dispersion for RNA-seq data with DESeq2. Genome Biol. 15, 1–21.

McInnes, L., Healy, J., and Melville, J. (2018). UMAP: Uniform Manifold Approximation and Projection for Dimension Reduction.

McVey, M., and Lee, S.E. (2008). MMEJ repair of double-strand breaks (director’s cut): deleted sequences and alternative endings. Trends Genet. 24, 529–538.

Mercier, F.E., Sykes, D.B., and Scadden, D.T. (2016). Single targeted exon mutation creates a true congenic mouse for competitive hematopoietic stem cell transplantation: The C57BL/6-CD45.1STEM mouse. Stem Cell Reports 6, 985–992.

Miqueu, P., Guillet, M., Degauque, N., Doré, J.C., Soulillou, J.P., and Brouard, S. (2007). Statistical analysis of CDR3 length distributions for the assessment of T and B cell repertoire biases. Mol. Immunol. 44, 1057–1064.

Mohrin, M., Bourke, E., Alexander, D., Warr, M.R., Barry-Holson, K., Le Beau, M.M., Morrison, C.G., and Passegué, E. (2010). Hematopoietic Stem Cell Quiescence Promotes Error-Prone DNA Repair and Mutagenesis. Cell Stem Cell 7, 174–185.

Naldini, L. (2019). Genetic engineering of hematopoiesis: current stage of clinical translation and future perspectives. EMBO Mol. Med. 11, 1–12.

Noda, S., Horiguchi, K., Ichikawa, H., and Miyoshi, H. (2008). Repopulating Activity of Ex Vivo-Expanded Murine Hematopoietic Stem Cells Resides in the CD48 − c-Kit + Sca-1 + Lineage Marker − Cell Population . Stem Cells 26, 646–655.

Obata, T., Suzuki, Y., Ogawa, N., Kurimoto, I., Yamamoto, H., Furuno, T., Sasaki, T., and Tanaka, M. (2014). Improvement of the Antitumor Activity of Poorly Soluble Sapacitabine (CS-682) by Using Soluplus ® as a Surfactant. 37, 802–807.

Pannetier, C., Cochet, M., Darche, S., Casrouge, A., Zoller, M., and Kourilsky, P. (1993). The sizes of the CDR3 hypervariable regions of the murine T-cell receptor β chains vary as a function of the recombined germ-line segments. Proc. Natl. Acad. Sci. U. S. A. 90, 4319–4323.

R Core Team (2020). R: A language and environment for statistical computing.

Rahman, S.H., Kuehle, J., Reimann, C., Mlambo, T., Alzubi, J., Maeder, M.L., Riedel, H., Fisch, P., Cantz, T., Rudolph, C., et al. (2015). Rescue of DNA-PK Signaling and T-Cell Differentiation by Targeted Genome Editing in a prkdc Deficient iPSC Disease Model. PLoS Genet. 11, 1–21.

De Ravin, S.S., Li, L., Wu, X., Choi, U., Allen, C., Koontz, S., Lee, J., Theobald-Whiting, N., Chu, J., Garofalo, M., et al. (2017). CRISPR-Cas9 gene repair of hematopoietic stem cells from patients with X-linked chronic granulomatous disease. Sci. Transl. Med. 9.

Riesenberg, S., Chintalapati, M., Macak, D., Kanis, P., and Maricic, T. (2019). Simultaneous precise editing of multiple genes in human cells. Nucleic Acids Res.

Schiroli, G., Ferrari, S., Conway, A., Jacob, A., Capo, V., Albano, L., Plati, T., Castiello, M.C., Sanvito, F., Gennery, A.R., et al. (2017). Preclinical modeling highlights the therapeutic potential of hematopoietic stem cell gene editing for correction of SCID-X1. Sci. Transl. Med. 9.

Schiroli, G., Conti, A., Ferrari, S., della Volpe, L., Jacob, A., Albano, L., Beretta, S., Calabria, A., Vavassori, V., Gasparini, P., et al. (2019). Precise Gene Editing Preserves Hematopoietic Stem Cell Function following Transient p53-Mediated DNA Damage Response. Cell Stem Cell 0, 1–15.

Sharma, R., Dever, D.P., Lee, C.M., Azizi, A., Pan, Y., Camarena, J., Köhnke, T., Bao, G., Porteus, M.H., and Majeti, R. (2021). The TRACE-Seq method tracks recombination alleles and identifies clonal reconstitution dynamics of gene targeted human hematopoietic stem cells. Nat. Commun. 12, 1–12.

Sheridan, C. (2021). CRISPR therapies march into clinic, but genotoxicity concerns linger. Nat. Biotechnol. 39, 897–899.

Tomellini, E., Fares, I., Lehnertz, B., Chagraoui, J., Mayotte, N., MacRae, T., Bordeleau, M.È., Corneau, S., Bisaillon, R., and Sauvageau, G. (2019). Integrin-α3 Is a Functional Marker of Ex Vivo Expanded α Human Long-Term Hematopoietic Stem Cells. Cell Rep. 28, 1063–1073.e5.

Tsai, S.Q., and Joung, J.K. (2016). Defining and improving the genome-wide specificities of CRISPR-Cas9 nucleases. Nat. Rev. Genet. 17, 300–312.

Vazquez, S.E., Inlay, M.A., and Serwold, T. (2015). CD201 and CD27 identify hematopoietic stem and progenitor cells across multiple murine strains independently of Kit and Sca-1. Exp. Hematol. 43, 578–585.

Wilkinson, A.C., Ishida, R., Kikuchi, M., Sudo, K., Morita, M., Crisostomo, R.V., Yamamoto, R., Loh, K.M., Nakamura, Y., Watanabe, M., et al. (2019). Long-term ex vivo haematopoietic-stem-cell expansion allows nonconditioned transplantation. Nature 571, 117–121.

Wilkinson, A.C., Igarashi, K.J., and Nakauchi, H. (2020). Haematopoietic stem cell self-renewal in vivo and ex vivo. Nat. Rev. Genet. 21, 541–554.

Wilkinson, A.C., Dever, D.P., Baik, R., Camarena, J., Hsu, I., Charlesworth, C.T., Morita, C., Nakauchi, H., and Porteus, M.H. (2021). Cas9-AAV6 gene correction of beta-globin in autologous HSCs improves sickle cell disease erythropoiesis in mice. Nat. Commun. 12, 686.

Wu, T., Hu, E., Xu, S., Chen, M., Guo, P., Dai, Z., Feng, T., Zhou, L., Tang, W., Zhan, L., et al. (2021). clusterProfiler 4.0: A universal enrichment tool for interpreting omics data. Innov. 2, 100141.

Zebedee, S.L., Barritt, D.S., and Raschke, W.C. (1991). Comparison of Mouse Ly5a and Ly5h Leucocyte Common Antigen Alleles. Dev. Immunol. 1, 243–254.

