## Supplemental Tables S1-S2 for "A Single Cell Cloning Platform for Gene Edited Functional Murine Hematopoietic Stem Cells"

**Supplemental Table 1. Candidate off-target sites, related to Fig. 1**

Candidate off-target sites of the guide RNA targeting *Prkdc* were identified using the COSMID algorithm. Mismatches and permutations in the seed region are indicated in red. The first row shows the seed sequence and PAM of the sgRNA used (Prkdc_gRNA1).

| **#** | **Hit** | **Permutation** | **Mismatches** | **Position (GRCm38)** | **strand** | **Gene** |
| --- | --- | --- | --- | --- | --- | --- |
|  | CTTACCAAGTTATAACAGCTNGG | - | - | - | - | Prkdc_gRNA1 |
| **1** | **A**TTAC**TT**AGTTATAACAGCTGGG | No indel | 3 | Chr5:57647418-57647440 | - | 4932441J04Rik |
| **2** | **C**TTA**A**AAAGT**C**ATAACAGCTGGG | No indel | 3 | Chr5:21558299-21558321 | - | Fbxl13; Lrrc17 |
| **3** | CTTA**-T**AA**C**TTATAACAGCTAGG | Del 15, or Del 16 | 2 | Chr14:96347342-96347363 | - | Klhl1 |
| **4** | CTTA**A**CAA**-**TT**C**TAACAGCTGGG | Del 12 | 2 | Chr8:73726599-73726620 | + | No known gene |
| **5** | CTTA**G**CAA**-**TTTTAACAGCTCGG | Del 12 | 2 | Chr1:24130829-24130850 | - | No known gene |
| **6** | CTT**C**CCAAG**-**TA**G**AACAGCTGGG | Del 10, or Del 11 | 2 | Chr4:54112620-54112641 | + | No known gene |
| **7** | CTT**C**CCAAG**-**TAT**T**ACAGCTTGG | Del 10, or Del 11 | 2 | Chr2:140296967-140296988 | + | Sel1l2 |

**Supplemental Table 2. Oligonucleotides used in this study, related to STAR Methods**

Sequences of DNA and RNA oligos used in this work.

| **Sequence** | **Source** | **Identifier** |
| --- | --- | --- |
| **Synthetic guide RNA (sgRNA) (mN*: Phosphorothioated 2'-O-methyl RNA base)** |  |  |
| mC*mU*mU*ACCAAGUUAUAACAGCUGUUUUAGAGCUAGAAAUAGCAAGUUAAAAUAAGGCUAGUCCGUUAUCAACUUGAAAAAGUGGCACCGAGUCGGUGCmU*mU*mU*U | This study | Prkdc_gRNA1 |
| mA*mU*mA*CUUCAAUUUGUUUGGAGGUUUUAGAGCUAGAAAUAGCAAGUUAAAAUAAGGCUAGUCCGUUAUCAACUUGAAAAAGUGGCACCGAGUCGGUGCmU*mU*mU*U | This study | Ptprc_gRNA1 |
| **Single-strand oligodeoxynucleotide (ssODN) HDR templates (N*: Phosphorothioated DNA base)** |  |  |
| G*G*A*TTCAAGAAATAAATGTAACGGAAAAGAATTGGTATCCACAACATAAAATACGCTATGCTAAGAGGAAGTTAGCAGGTGCCAATCCAGCTGTTATAACTTGGTAAGACTTGTGAATGCAGAA*T*C*A | This study | Prkdc_HDR_ssODN_asym |
| T*G*C*CCAGCATCGTACCTGGCTCACAGTGGAGTACATATGAAATATTGTCACTGTTGCATTTTCTGAAATCAAGGTTTTCTGTCTTCCATTCCAAACAAATGGAAGTATTAGCCTTTTCTTTTGG*T*G*T | This study | Ptprc_HDR_ssODN_asym |
| **PCR and sequencing primers (FAM-: 5' 6-carboxyfluorescein label)** |  |  |
| CTTTGTTTTAGGGTCATTACTTGGT | This study | Prkdc_inner_F |
| TGCTCAGAACTGAAGTCTAAGGT | This study | Prkdc_inner_R |
| CAATTATCCAGACTATCCCCGAAA | This study | Prkdc_outer_F |
| TCTTGCCTACACCCTGTAAAGC | This study | Prkdc_outer_R |
| TCCCTCCTAGAAGCACTTGTT | This study | Ptprc-e7-F |
| CCCCTAGCGAAATCTCCTGC | This study | Ptprc-e7-R |
| AGCGGAAGGGCAACTTTACTA | This study | OT_01_outer_F |
| GATCTCATACACAGACAGGGAAG | This study | OT_01_inner_F |
| GCATACGCCATTCTGCTCAC | This study | OT_01_inner_R_SEQ^a^ |
| TGCCTGTATGGTAAGCACCC | This study | OT_02_inner_F_SEQ^a^ |
| TTAGTGGCAGGCATGCTTCA | This study | OT_02_outer_R |
| AGTCACCATTCGTGTGTCCC | This study | OT_02_inner_R |
| TGGATCTTGTTCATCTGGGGC | This study | OT_03_inner_F_SEQ^a^ |
| GCCTGAGGGACTCAGTATTGT | This study | OT_03_outer_R |
| AGATTTCAAGATGTCCTTACGA | This study | OT_03_inner_R |
| TATTAGCACCTACACCAATGCT | This study | OT_04_outer_F |
| GTGGCACAAAGAAAGATGTATGG | This study | OT_04_inner_F |
| TGCAGTCCTTGTAAGAGGGT | This study | OT_04_inner_R_SEQ^a^ |
| GATGCAGCTGAGAGACTCGT | This study | OT_05_inner_F_SEQ^a^ |
| CACTCCCTGTGTGTTTGTTTC | This study | OT_05_inner_R |
| AGTAGGTCTTTGTAGGCACGC | This study | OT_05_outer_R |
| CGACAACACTCTGACTCCCATA | This study | OT_06_outer_F |
| CCTTTCCCTTGGGTACTTCTTG | This study | OT_06_inner_F |
| TGCAAATGACCGGAAATCTGTAAA | This study | OT_06_inner_R |
| GCCCTCAAGATTTTGCTGTCAAG | This study | OT_06_SEQ^a^ |
| TACCTTCTCACAAGCAGGGAGG | This study | OT_07_outer_F |
| GAGGAGCCTTATGGAAGAGTTG | This study | OT_07_inner_F |
| TTAAGGCATTCCTGTCTGCCA | This study | OT_07_inner_R_SEQ^a^ |
| CTGAATGCCCAGACAGCTCCAAGC | Ahmed et al., 2009 | TCR-Vb1 |
| CATTACTCATATGTCGCTGAC | Ahmed et al., 2009 | TCR-Vb8.1 |
| TGCTGGCAACCTTCGAATAGGA | Ahmed et al., 2009 | TCR-Vb8.3 |
| TCTCTCTACATTGGCTCTGCAGGC | Ahmed et al., 2009 | TCR-Vb9 |
| CTTGGGTGGAGTCACATTTCT | Ahmed et al., 2009 | TCR-Cb |
| FAM-CTTGGGTGGAGTCACATTTCT | Ahmed et al., 2009 | TCR-Cb-FAM |
| ^a^SEQ denoted off-target PCR primers were also used as sequencing primers |  |  |
