## Supplementary material for "A Single Cell Cloning Platform for Gene Edited Functional Murine Hematopoietic Stem Cells": Key Reagents

| **REAGENT or RESOURCE** | **SOURCE** | **IDENTIFIER** | |
| --- | --- | --- | --- |
| **Antibodies** | | | |
| anti-mouse Ckit-APC (2B8) | Thermo Fisher Scientific | Cat#17-1171-83; RRID:AB_469431 | |
| anti-mouse CD4-APC (RM4-5) | Thermo Fisher Scientific | Cat#17-0042-83; RRID:AB_469324 | |
| anti-mouse CD8a-APC (53-6.7) | Thermo Fisher Scientific | Cat#17-0081-83; RRID:AB_469336 | |
| anti-mouse CD48-APC (HM48-1) | Thermo Fisher Scientific | Cat#17-0481-82, RRID:AB_469408 | |
| anti-mouse CD135-APC (A2F10) | Thermo Fisher Scientific | Cat#17-1351-82, RRID:AB_10717261 | |
| anti-mouse CD45.1-APC/Cy7 (A20) | Tonbo Biosciences | Cat#25-0453; RRID:AB_2621629 | |
| anti-mouse Sca-1-APC/Cy7 (D7) | BioLegend | Cat#108126; RRID:AB_10645327 | |
| anti-mouse CD45R(B220)-APC/eFluor780 (RA3-6B2) | Thermo Fisher Scientific | Cat#47-0452-82; RRID:AB_1518810 | |
| anti-mouse Ckit-APC/H7 (2B8) | BD Biosciences | Cat#560185, RRID:AB_1645231 | |
| anti-mouse CD4-Biotin (RM4-5) | Thermo Fisher Scientific | Cat#13-0042-85; RRID:AB_466330 | |
| anti-mouse CD8a-Biotin (53-6.7) | Thermo Fisher Scientific | Cat#13-0081-85; RRID:AB_466347 | |
| anti-mouse CD45R(B220)-Biotin (RA3-6B2) | Thermo Fisher Scientific | Cat#13-0452-82; RRID:AB_466449 | |
| anti-mouse TER119-Biotin (TER119) | Thermo Fisher Scientific | Cat#13-5921-85; RRID:AB_466798 | |
| anti-mouse Gr1-Biotin (RB6-8C5) | Thermo Fisher Scientific | Cat#13-5931-85-85; RRID:AB_466801 | |
| anti-mouse CD11b-Biotin (M1/70) | Thermo Fisher Scientific | Cat#13-0112-85; RRID:AB_466360 | |
| anti-mouse CD127-Biotin (A7R34) | Thermo Fisher Scientific | Cat#13-1271-85; RRID:AB_466589 | |
| anti-mouse CD45.2-BV421 (104) | Thermo Fisher Scientific | Cat#48-0454-82; RRID:AB_11042125 | |
| anti-mouse CD34-FITC (RAM34) | Thermo Fisher Scientific | Cat#11-0341-85; RRID:AB_465022 | |
| anti-mouse CD45.2-FITC (104) | BioLegend | Cat#109806; RRID:AB_313443 | |
| anti-mouse CD48-FITC (HM48-1) | BioLegend | Cat#103404, RRID:AB_313019 | |
| anti-mouse CD4-FITC (RM4-5) | Tonbo Biosciences | Cat#35-0042, RRID:AB_2621666 | |
| anti-mouse CD8a-FITC (53-6.7) | Thermo Fisher Scientific | Cat#11-0081-85, RRID:AB_464916 | |
| anti-mouse CD45R(B220)-FITC (RA3-6B2) | Thermo Fisher Scientific | Cat#11-0452-85, RRID:AB_465055 | |
| anti-mouse TER119-FITC (TER119) | Thermo Fisher Scientific | Cat#11-5921-82, RRID:AB_465311 | |
| anti-mouse Gr1-FITC (RB6-8C5) | Thermo Fisher Scientific | Cat#11-5931-82, RRID:AB_465314 | |
| anti-mouse CD11b-FITC (M1/70) | Thermo Fisher Scientific | Cat# 11-0112-82, RRID:AB_464935 | |
| anti-mouse CD4-PB (RM4-5) | BioLegend | Cat#100531; RRID:AB_493374 | |
| anti-mouse CD8a-PB (53-6.7) | BD Biosciences | Cat#558106; RRID:AB_397029 | |
| anti-mouse CD45R(B220)-PB (RA3-6B2) | BD Biosciences | Cat#558108; RRID:AB_397031 | |
| anti-mouse TER119-PB (TER119) | BioLegend | Cat#116231; RRID:AB_2149212 | |
| anti-mouse Gr1-PB (RB6-8C5) | BioLegend | Cat#108430; RRID:AB_893556 | |
| anti-mouse CD11b-PB (M1/70) | BioLegend | Cat#101224; RRID:AB_755986 | |
| anti-mouse Sca-1-PE (D7) | BD Biosciences | Cat#12-5981-83; RRID:AB_466087 | |
| anti-mouse CD201-PE (eBio1560) | Thermo Fisher Scientific | Cat#12-2012-82; RRID:AB_914317 | |
| anti-mouse Gr1-PE (RB6-8C5) | Thermo Fisher Scientific | Cat#12-5931-83; RRID:AB_466046 | |
| anti-mouse CD11b-PE (M1/70) | BD Biosciences | Cat#557397; RRID:AB_396680 | |
| anti-mouse CD105-PE (MJ7/18) | Thermo Fisher Scientific | Cat#12-1051-82, RRID:AB_657524 | |
| anti-mouse CD150-PE/Cy7 (TC15-12F12.2) | BioLegend | Cat#115913; RRID:AB_439796 | |
| anti-mouse CD45.1-PE/Cy7 (A20) | Tombo Biosciences | Cat#60-0453; RRID:AB_2621850 | |
| anti-mouse CD8a-PE/Cy7 (53-6.7) | BioLegend | Cat#100722; RRID:AB_312761 | |
| anti-mouse Sca-1-BV605 (D7) | BioLegend | Cat#108133, RRID:AB_2562275 | |
| Streptavidin-APC/eFluor780 | Thermo Fisher Scientific | Cat#47-4317-82; RRID:AB_10366688 | |
| Streptavidin-BV421 | BioLegend | Cat#405225 | |
| **Chemicals, Peptides, and Recombinant Proteins** | | | |
| Polyvinyl alcohol (PVA), 87-90% hydrolyzed | Sigma | Cat#P8136; CAS 9002-89-5 | |
| Soluplus | BASF | CAS 402932-23-4 | |
| Recombinant Murine TPO | Peprotech | Cat#315-14; P40226 | |
| Recombinant Murine SCF | Peprotech | Cat#250-03; P20826 | |
| Insulin-Transferrin-Selenium | Thermo Fisher Scientific | Cat#41400045 | |
| Nitroiodophenyl (NIP)-conjugated OVA (NIPOVA) | Biosearch Technologies | Cat# N-5041-10 | |
| **Deposited data** | | | |
| RNASeq data (normalized counts) | This paper | | https://doi.org/10.5061/dryad.m905qfv2f (not yet activated) |
| Raw and analyzed data | This paper | | https://doi.org/10.5061/dryad.m905qfv2f (not yet activated) |
| **Experimental Models: Cell Lines** | | | |
| Human: A549 (lung adenocarcinoma) | ATCC | Cat#CCL-185, RRID:CVCL_0023 | |
| **Experimental Models: Organisms/Strains** | | | |
| Mouse: CB17/SCID: C.B-17/Icr-^scid/scid^Jcl | Clea | RRID:IMSR_JCL:JCL:mID-0003 | |
| Mouse: CB17/WT: C.B-17/Icr-^+/+^Jcl | Clea | RRID:IMSR_JCL:JCL:mID-0004 | |
| Mouse: CD45.2^+^ C57BL/6: C57BL/6NCrSlc | SLC | RRID:MGI:5295404 | |
| Mouse: CD45.1^+^ C57BL/6: B6.SJL-Ptprc^a^ Pepc^b^/BoyJ | Sankyo Labo | RRID:IMSR_JAX:002014 | |
| **Oligonucleotides** | | | |
|  | See Table S2 for list of oligonucleotides |  | |
| **Software and Algorithms** | | | |
| FlowJo version 10 | BD | https://www.flowjo.com; SCR_008520 | |
| IndexSort version 0.1.6 (FlowJo plugin) | Freier, 2020 | https://www.flowjo.com | |
| UMAP version 3.1 (FlowJo plugin) | McInnes et al.,2018 | https://arxiv.org/abs/1802.03426 | |
| Prism version 9.1 | Graphpad | https://www.graphpad.com; SCR_002798 | |
| R version 4.0.0 | R Foundation | https://www.r-project.org; SCR_001905 | |
| ggplot2 version 3.3.5 | Wickham 2016 | https://ggplot2.tidyverse.org; SCR_014601 | |
| DESeq2 version 1.32.0 | Love et al., 2014 | SCR_015687 | |
| clusterProfiler version 4.0.2 | Wu et al., 2021 | SCR_016884 | |
| ComplexHeatmap version 2.8.0 | Gu et al., 2016 | SCR_017270 | |
| Inference of CRISPR edits (ICE) | Synthego | https://ice.synthego.com/ | |
| Tracking of Indels by Decomposition (TIDE) | Brinkman et al., 2014 | http://shinyapps.datacurators.nl/tide/ | |
